# SonicParanoid2: fast, accurate, and comprehensive orthology inference with machine learning and language models

**DOI:** 10.1101/2023.05.14.540736

**Authors:** Salvatore Cosentino, Wataru Iwasaki

**Affiliations:** Department of Integrated Biosciences, Graduate School of Frontier Sciences, the University of Tokyo, Kashiwa, Japan; Department of Biological Sciences, Graduate School of Science, the University of Tokyo, Bunkyo-ku, Japan; Department of Computational Biology and Medical Sciences, Graduate School of Frontier Sciences, the University of Tokyo, Kashiwa, Japan; Atmosphere and Ocean Research Institute, the University of Tokyo, Kashiwa, Japan; Institute for Quantitative Biosciences, the University of Tokyo, Bunkyo-ku, Japan; Collaborative Research Institute for Innovative Microbiology, the University of Tokyo, Bunkyo-ku, Japan

**Keywords:** Orthology inference, Machine learning, Language model, Genome evolution

## Abstract

Accurate inference of orthologous genes constitutes a prerequisite for comparative and evolutionary genomics. SonicParanoid is one of the fastest tools for orthology inference; however, its scalability and accuracy have been hampered by time-consuming all-versus-all alignments and the existence of proteins with complex domain architectures. Here, we present a substantial update of Sonicparanoid, where a gradient boosting predictor halves the execution time and a language model doubles the recall. Application to empirical large-scale and standardized benchmark datasets showed that SonicParanoid2 is up to 18X faster than comparable methods and also the most accurate. SonicParanoid2 is available at https://gitlab.com/salvo981/sonicparanoid2

## Background

The accurate inference of orthologous genes originating from speciation events is crucial in various areas of genomics and evolutionary biology [1]. Many tools and resources have been developed to identify orthologous relationships among multiple proteomes. They are classified into graph [2–6] and tree based [7–9], which do not include tools that integrate both approaches (hybrid) or integrate publicly available resources to perform their predictions [10,11]. Graph-based tools infer orthologs by calculating all-versus-all pairwise similarity scores and using the bidirectional-best-hit (BBH) method or its derivatives [12]. In tree-based methods, orthologs are identified by analyzing phylogenetic trees, which allows the identification of speciation and duplication events. Tree-based tools are more computationally demanding and are typically available to users only via databases or web services. A comparative study showed no significant differences in the ability of the two approaches in inferring orthologs [13].

The steep decrease in sequencing costs and the associated increase in genomic and metagenomic data [14] challenge the scalability of orthology inference tools. Inferring orthologs *de novo* for a few hundred proteomes using graph-based methods on high-performance computing servers may require days to weeks, even when local alignment tools, such as MMseqs2 [15] and Diamond [16], are used as faster alternatives to BLAST [17].

In addition to the scalability problem, graph-based methods tend to miss orthologs in duplication-rich proteomes [18] (e.g., plant proteomes) and in proteins with complex domain arrangements (architectures) originating from fusion and fission events [19,20]. For example, BBH-based methods fail to identify human kinase orthologs detected using a domain-based orthology inference method [21].

Machine learning (ML), particularly methods borrowed from natural language processing (NLP), has been extensively used in genomics [22]. Currently, deep learning and language models are used in multiple areas of genomics [23], including sequence assembly and binning [24,25] as well as protein folding [26,27]. ML is gradually being adopted for orthology inference [28,29]; however, the sensitivity, scalability, and usability of ML-based methods can be further improved. Herein, we report a major update to SonicParanoid [30], which is one of the fastest *de novo* orthology inference tools [29,31]. The update uses two ML methods, AdaBoost [32] and Doc2Vec [33], to deliver a substantially faster, more accurate, and more comprehensive orthology inference.

## Results

### SonicParanoid2 as the fastest and most accurate orthology inference method

For a set of *N* proteomes, SonicParanoid2 performs *de novo* orthology inference using a novel graph-based algorithm that halves the execution time with an AdaBoost classifier and avoiding unnecessary alignments (Figure 1a). Furthermore, for a more comprehensive identification of orthologs, SonicParanoid2 conducts domain-based orthology inference using Doc2Vec neural network models (Figure 1b). The orthologs predicted by these two algorithms are merged (Figure 1c) and input into the Markov cluster algorithm [34] (MCL) to infer the ortholog groups [2] (OGs) for the *N* input proteomes. SonicParanoid2 can be executed using three predefined modes (fast, default, and sensitive), in which different local alignment tools and settings are used (Supplementary Table S1).

**Figure 1:**
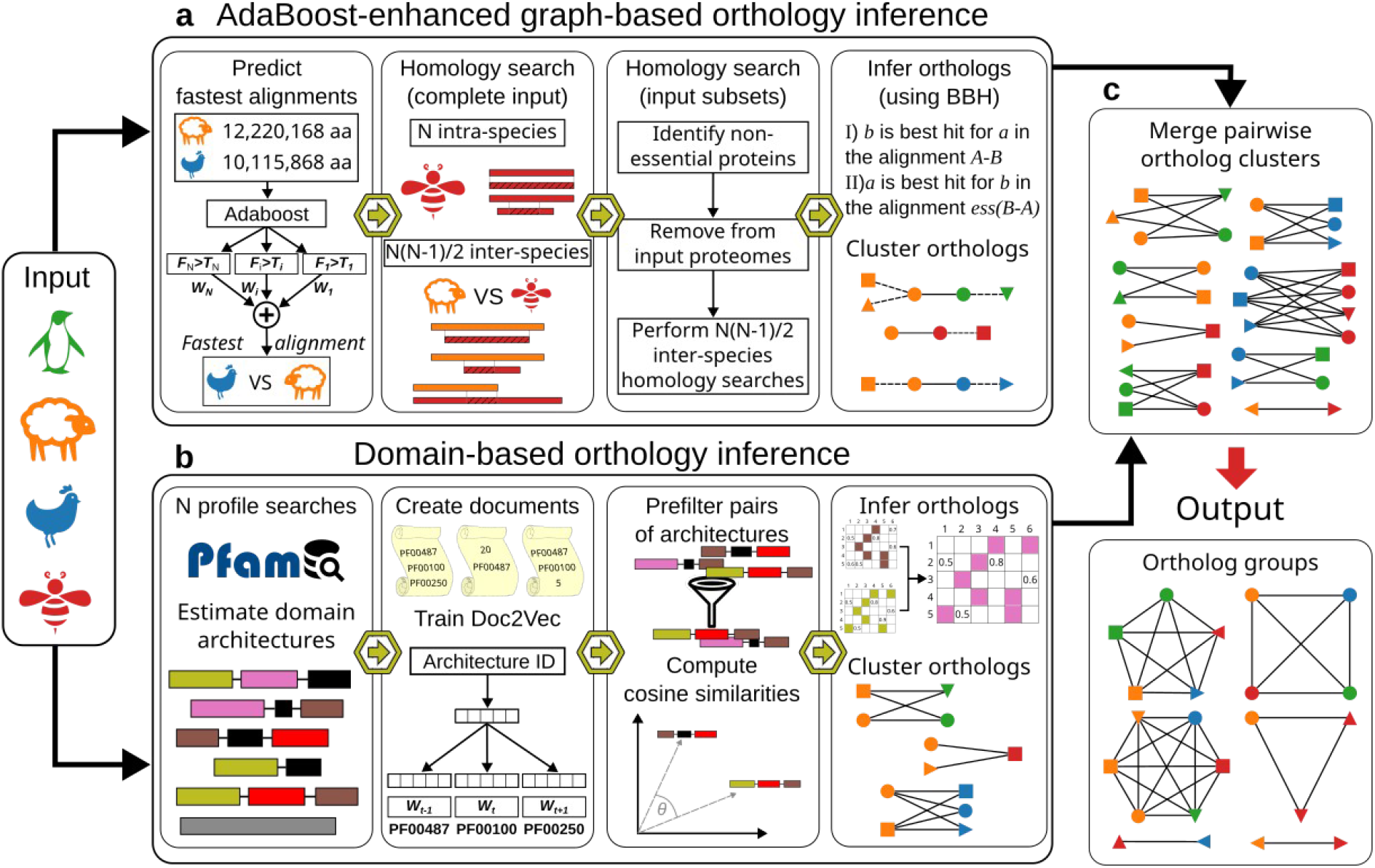
Overview of SonicParanoid2. **a**, Graph-based orthology inference pipeline using a novel ML-based approach to substantially reduce execution time of homology searches; **b**, Domain-based orthology inference pipeline that compares domain architectures using methods from NLP; **c**, Pairwise orthologous tables predicted using the two pipelines combined to generate ortholog graphs from which the output ortholog groups are inferred.

The effect of domain-based orthology on the number of predicted orthologs and the total execution time differs depending on the input dataset and settings for SonicParanoid2. For example, on a dataset provided by the Quest for Orthologs [35,36] (QfO)(Supplementary Figure S1a), the number of predicted orthologs increased by one-third at the default settings, and the total execution time showed a similar increase (29.17% (10.70’)) (Table 1). The increase in the total execution time becomes relatively negligible for larger datasets and at higher sensitivity settings because of the high scalability of the domain-based pipeline (Supplementary Figure S2). For example, on a dataset with 2,000 metagenome-assembled genomes (MAGs) (Supplementary Figure S1b), the increase in execution time was 10% and 24% for the default and fast modes, respectively, whereas one-third more orthologs were predicted regardless of the settings (Table 2).

**Table 1:** Execution times of SonicParanoid2 and effects of including domain-based orthology. Results were obtained using the QfO dataset as input. Execution time columns show the total execution time and its increase due to the execution of the domain-based pipeline. Increase in predicted orthologs provided by the domain-aware pipeline is up to one-third (columns of predicted ortholog pairs). Last two columns show speedup folds relative to the execution times of Broccoli and OrthoFinder (both at default settings).

| Mode | Execution time |  | Predicted ortholog pairs |  | Speed-up on other methods |  |
| --- | --- | --- | --- | --- | --- | --- |
|  | Total (hours) | Increase (%) | Count (million) | Increase (%) | Broccoli (folds) | OrthoFinder (folds) |
| Default | 0.86 | 29.17 | 15.28 | 27.39 | 3.23 | 11.89 |
| Fast | 0.56 | 49.16 | 14.64 | 29.79 | 4.92 | 18.10 |
| Sensitive | 4.36 | 4.45 | 19.27 | 23.14 | -1.57 | 2.34 |

**Table 2:** Effects of including domain-oriented orthology when processing a set of 2,000 MAGs. Execution time columns show the total execution time and its increase due to the inclusion of the domain-based pipeline. The increase in execution time is low if we consider the magnitude of the input dataset, whereas the number of predicted orthologs increased by more than one-third.

| Mode | Execution time |  | Predicted ortholog pairs |  |
| --- | --- | --- | --- | --- |
|  | Total (hours) | Increase (%) | Count (million) | Increase (%) |
| Default | 37.99 | 10.40 | 1817.40 | 30.66 |
| Fast | 24.33 | 23.51 | 1694.88 | 37.39 |

We compared the execution time of SonicParanoid2 with those of OrthoFinder [10] and Broccoli [29], which are among the fastest tools available for *de novo* orthology inference. Broccoli reduces the burden of all-versus-all alignments through *k*-mer clustering and uses label propagation instead of the typically used MCL to infer OGs. Multiple studies demonstrated that these three tools are equally accurate for inferring OGs [31,37,38].

On the QfO dataset, SonicParanoid2 was 18X and 5X faster than OrthoFinder and Broccoli, respectively, in the fast mode (Table 1). At the default settings, SonicParanoid2 was 12X and 3X faster, respectively. In the sensitive mode, SonicParanoid2 was approximately 1.5X slower than Broccoli. At these settings, MMseqs2 at the highest sensitivity was used (Supplementary Table S1), whereas Broccoli and OrthoFinder used Diamond which is faster. Orthologous relationships for the MAG dataset were inferred by SonicParanoid2 in 1.6 and 1 day in the default and fast modes, respectively (Table 2). Conversely, OrthoFinder and Broccoli failed to complete the orthology inference on the MAG dataset within eight days (i.e., more than 8X longer than SonicParanoid2).

The accuracy of SonicParanoid2 was evaluated using the QfO benchmark [36] and compared with 14 well-established methods, including a legacy version of SonicParanoid [30]. In 12 tests performed, methods close to or constituting the Pareto frontier were regarded as those that provided the best tradeoff between precision and recall. SonicParanoid2 was Pareto optimal in multiple tests, including the LUCA and bacterial species tree discordance tests (Figure 2a-c and Supplementary Figures S3 and S4). Moreover, SonicParanoid2 is the most accurate method based on the aggregate ranking of three classification methods provided by the QfO benchmarking service (Figure 2d and Supplementary Figures S5-S8).

**Figure 2:**
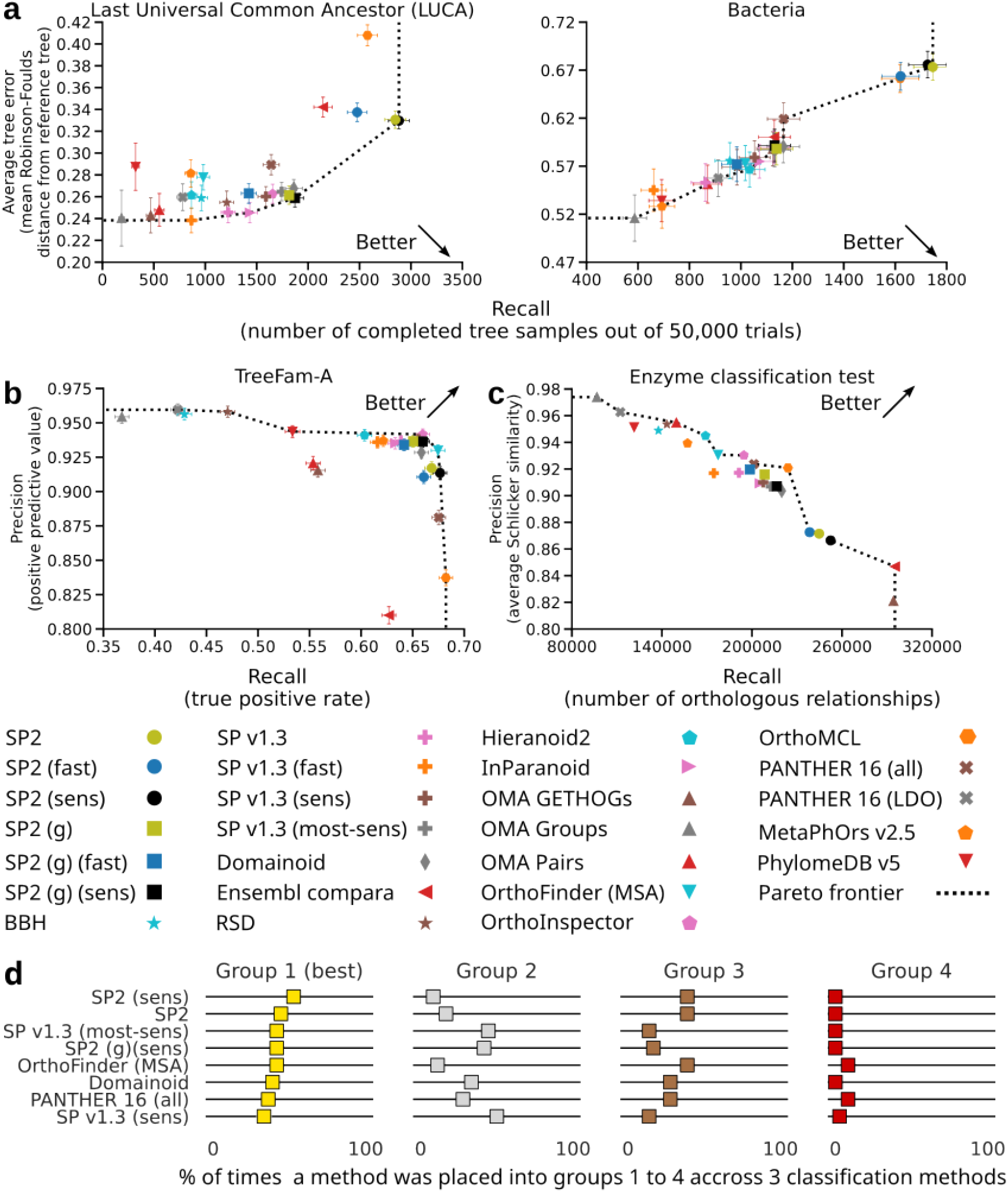
Accuracy of SonicParanoid2 compared with other 14 methods. **a-c**, Accuracies of SonicParanoid2 (SP2) using different settings on four among 12 QfO 2020 benchmark tests. Methods lying on Pareto frontier indicate the best balance between precision and recall. Results obtained using only the novel graph-based approach are shown (squares) for comparison with older versions of SonicParanoid (crosses). **a**, Species tree discordance tests on LUCA and bacteria datasets. **b**, Reference gene tree test based on TreeFam-A. **c**, Functional benchmark test based on Enzyme Commission (EC) numbers. **d**, Top eight methods according to the aggregate ranking from the three classification methods in the QfO benchmark.

### Substantially faster and scalable graph-based orthology inference using machine learning

Graph-based *de novo* orthology inference for *N* proteomes generally requires *N* intra-proteome and *N(N-1)* bidirectional inter-proteome all-versus-all alignments. For the bidirectional inter-proteome alignment of proteomes A and B, all proteins in A are queried against proteome B (denoted as alignment A-B), and all proteins in B are queried against proteome A (B-A). The key rationale behind the way in which SonicParanoid2 reduces the computational cost are: (1) The execution times of alignments A-B and B-A can substantially differ depending on the proteome size and evolutionary distance between A and B (Supplementary Figs. S9 and S10), and (2) query and target proteins that have no hits with bitscores above a predefined threshold in one of the two inter-proteome alignments cannot be predicted as orthologs, based on the definition of BBH (Equation 1). Accordingly, for a pair of proteomes A and B, SonicParanoid2 first predicts the faster alignment between A-B and B-A using an adaptive gradient boosting [32] (AdaBoost) binary classifier (Supplementary Table S2). If A-B is predicted to be faster, then SonicParanoid2 first conducts the alignment A-B and creates two subsets, ess(A) and ess(B), which include only proteins with alignment scores above the threshold (Equation 2). Subsequently, the alignment between ess(B) and ess(A) is performed instead of the expectedly slower alignment of B-A (Supplementary Fig. S11). Finally, the algorithm identifies the orthologs as shown in Equation 3. Henceforth, we will refer to the subsets of proteins generated by this graph-based algorithm as “essential subsets” and to the all-versus-all among these subsets as “essential alignments”.

We evaluated the execution time of the novel graph-based algorithm in SonicParanoid2 (Figure 1a) on the two proteome sets shown in Supplementary Figure S1 using different alignment tools and sensitivity settings (Supplementary Table S3). For the QfO dataset, the execution time for the all-versus-all alignments was reduced by 42% (Supplementary Table S4). Moreover, when only inter-proteome alignments were considered, the reduction in execution time was as high as 95% (Figure 3). A speedup was observed regardless of the alignment tool used (MMseqs2, Diamond, or BLAST). Nevertheless, the extent depended on both the alignment tool used and its sensitivity settings (Supplementary Table S4). Specifically, the proportions of sequences used in the second alignment were smaller for the proteome pairs of phylogenetically distant species (Supplementary Figure S12), thus resulting in higher speedups. For example, 97% of the input proteins was not used in the second alignment between *Leptospira interrogans* (bacteria) and *Giardia intestinalis* (protist), which consequently reduced the execution time substantially. Conversely, less than 2% of the original input was removed for the second alignment between closely related chimpanzee and gorilla proteomes.

**Figure 3:**
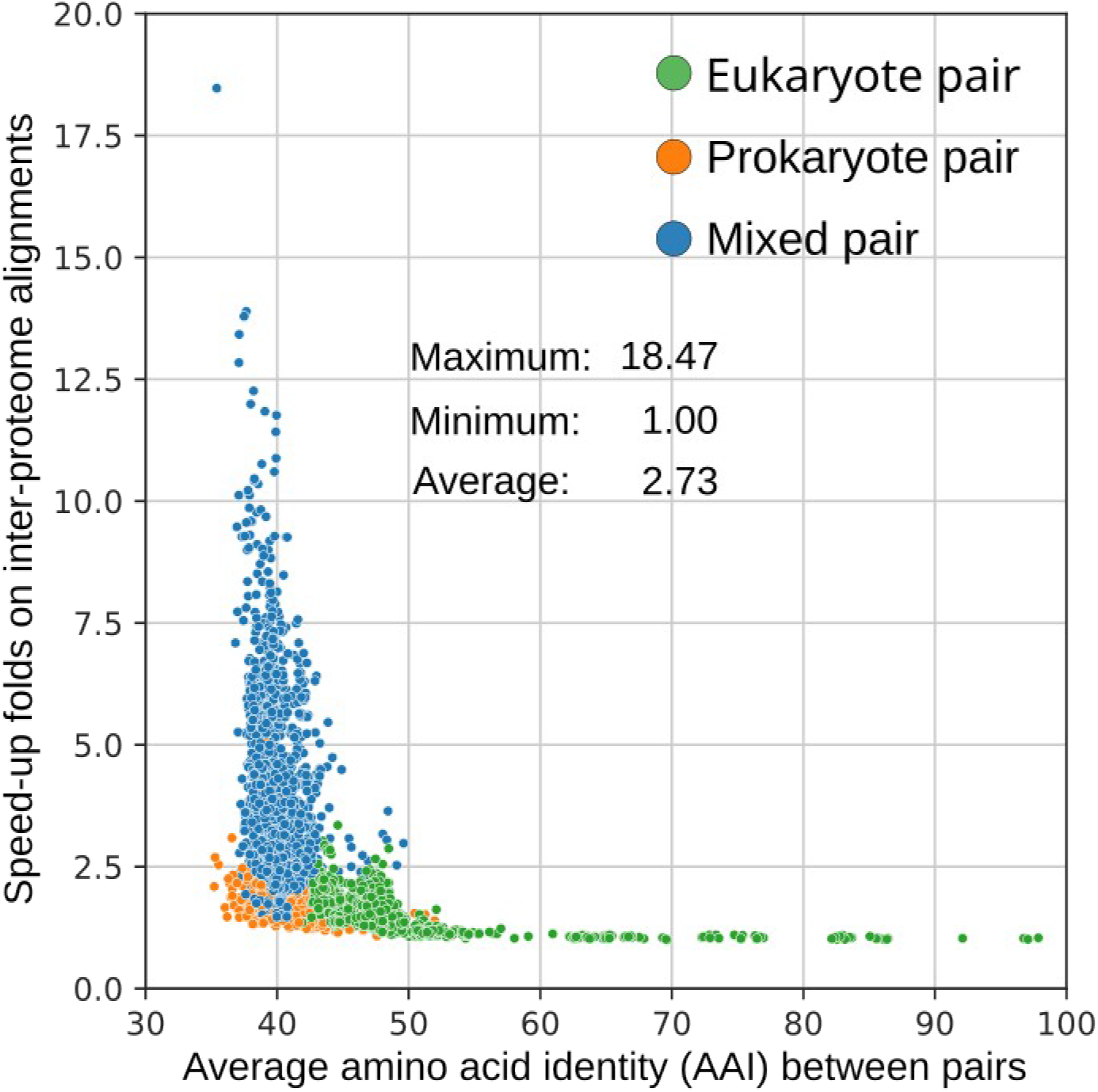
Speedup from essential alignments and evolutionary relatedness. For two proteomes A and B, the total execution time is computed as the sum of the execution times for the inter-proteome alignments A-B and B-A. The speedup (on the Y-axis) obtained for each bidirectional alignment was computed as the difference in total execution time required with and without using the essential alignments. The speedups obtained using the novel approach in SonicParanoid2 are inversely proportional to the evolutionary relatedness (expressed in terms of AAI) of the proteome pairs. The results above were obtained using the QfO dataset as input and MMseqs2 at the most sensitive settings.

The accuracy of the AdaBoost classifier differed depending on the alignment tool for which it inferred faster alignments, partly because it was trained for MMseqs2 (Supplementary Table S4). The necessity of the AdaBoost model was assessed by measuring the reduction in execution time when the slowest alignments were conducted first. This task was performed by inverting the predictions of the AdaBoost classifier. For example, when the classifier predicted A-B as the fastest alignment, we first aligned B-A. We observed that the amount of saved time was up to 70% less, which proved the importance of predicting the fastest inter-proteome alignments (Supplementary Table S5).

To evaluate the scalability of the graph-based algorithm of SonicParanoid2, we performed orthology inference on a large dataset comprising 2,000 MAGs from aquatic and terrestrial environments (Supplementary Figure S1b). The required four million all-versus-all alignments were performed in 56.48 and 23.74 hours using MMseqs2 and Diamond, respectively, which corresponded to execution times reduced by 16.18% and 24.99%, respectively (Supplementary Table S6). The MAG dataset contained many closely related microbial genomes, and the closely related proteomes therein limited the reduction in execution time (Supplementary Figures S13 and S12).

### Speed and accuracy of the AdaBoost-enhanced graph-based method

We compared the execution times of the AdaBoost-enhanced graph-based algorithm (Figure 1a) with those of Broccoli and OrthoFinder. In particular, we compared the execution times for the alignment-based orthology inference alone; hence, SonicParanoid2 was set to omit the domain-based orthology. SonicParanoid2 executed in the fast mode using Diamond (Supplementary Table S3) completed the execution in 0.38 hours for the 78-proteome QfO dataset, whereas OrthoFinder and Broccoli required 10.19 and 2.77 hours (7.3 and 26.9-folds slower), respectively (Supplementary Table S7). Subsequently, we tested OrthoFinder and Broccoli on the 2,000-proteome MAG dataset; however, both tools failed to complete the orthology inference within eight days, whereas SonicParanoid2 required one day (Supplementary Table S6).

To compare the AdaBoost-enhanced method with the conventional graph-based approach, which uses all the input sequences, we evaluated the prediction sets of 18 trials (Supplementary Table S4) using the QfO benchmark. The accuracies of the two graph-based approaches were highly similar, with no apparent adverse effect on the accuracy despite the substantial reduction in computational time (Supplementary Figures S14 and S15). This can better be seen in Supplementary figures S5-8 in which “SP v1.3 (most-sens)”, and “SP2 (g)(sens)” are ranked next to each other in three of the four rankings. These two methods used only the graph-based algorithm and the same settings for MMseqs (Supplementary Tables S1 and S3).

### Fast and scalable domain-based orthology inference using Doc2Vec

The graph-based approach based on BBH can miss orthologs of proteins with many duplications or those that have undergone domain fusion or fission [18,20,21] (Figure 4a and Supplementary Figure S16). Thus, in addition to the orthologs predicted using the graph-based approach, SonicParanoid2 infers orthologs at the domain level by comparing functional domain architectures using techniques typically used in natural language processing (Figures 1b and 4b).

**Figure 4:**
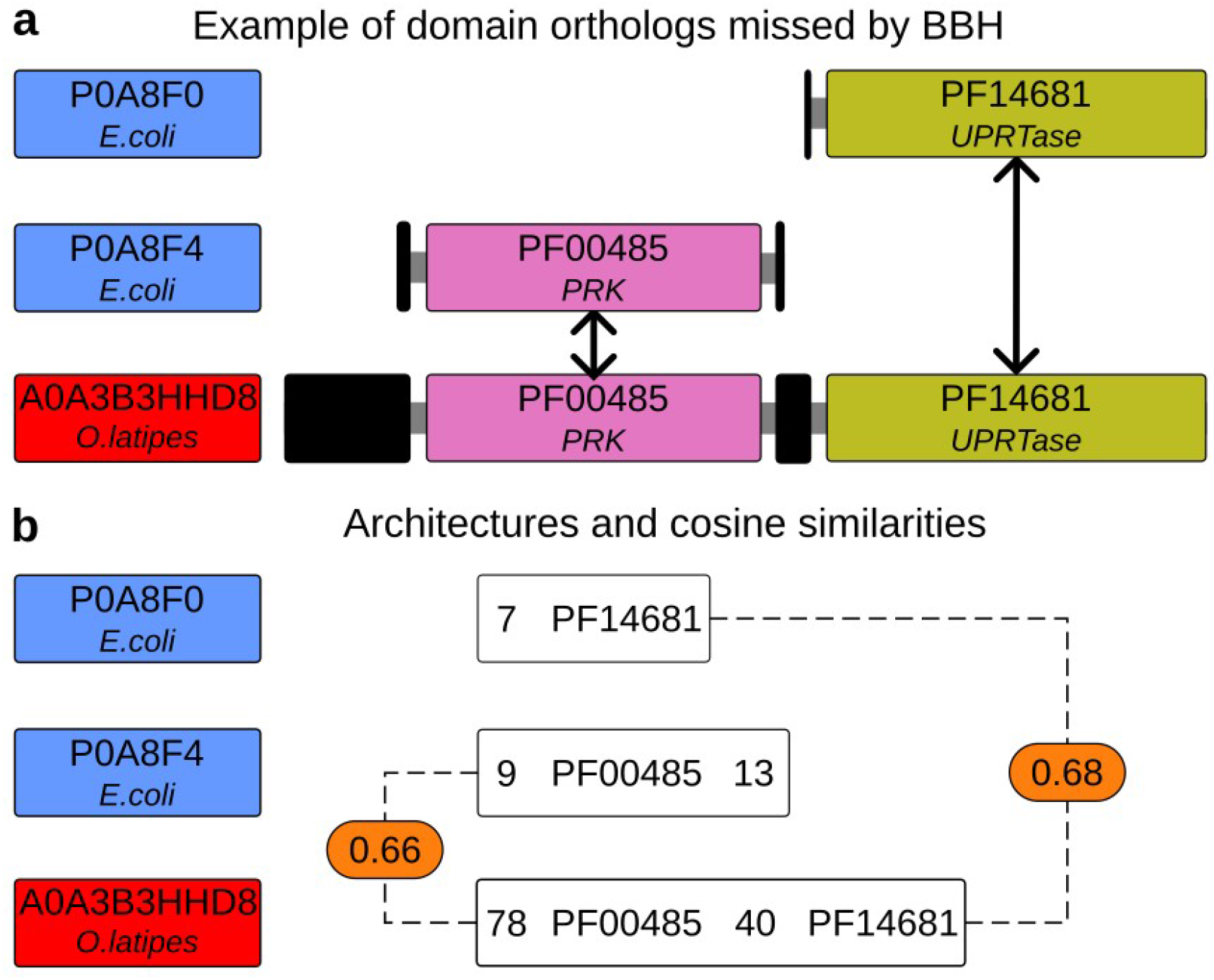
Recovery of orthologs missed by BBH via domain-based orthology inference. **a**, Protein A0A3B3HHD8 from *O. latipes* is a representative example of domain fusion resulting from two single-domain *E. coli* proteins. Pink and green boxes in **a** are functional domains (annotated with their Pfam ID and gene name), whereas black boxes represent inter-regions with no Pfam annotation in InterPro (as of January 2023). P0A8F4 and P0A8F0 are orthologous to A0A3B3HHD8 (black arrows), but the graph-based pipeline identifies only P0A8F4 as an ortholog. **b**, Using domain-aware orthology, SonicParanoid2 predicts P0A8F0 as an ortholog of A0A3B3HHD8. Text inside the white boxes represent the architectures for the proteins above as they appear in the training corpus for the Doc2Vec model. The rounded rectangles in orange show the cosine similarities assigned to the two pairs of domain architectures.

SonicParanoid2 uses fast profile searches on Pfam [39] to infer the domain architectures of the input proteins and converts them into “phrases”, where “words” are the annotated functional domains and the amino-acid lengths of the inter-domain regions (Figure 1b). The phrases are used as the training corpus for a Doc2Vec [33] neural network model, that associates a numerical vector with each domain architecture. For each inter-proteome pair, after a filtering step, the cosine similarities of the inter-proteome domain architecture pairs are computed. Pairs of architectures with cosine similarities above a predefined threshold are considered candidate orthologs.

The increase in total execution time due to the inclusion of the domain-based pipeline was negligible, considering the increase in the number of predicted orthologs (Table 1). Due to the high scalability of the domain-based pipeline (Supplementary Figure S2), the aforementioned smallness of the increase in execution time was more apparent when processing larger datasets. For example, for the MAG dataset, the increase in execution time was 10% and 24% for the default and fast modes, respectively, whereas the increase in the number of predicted orthologs was higher than one-third, regardless of the settings (Table 2).

The inclusion of the domain-based pipeline substantially increased the recall. For example, in the bacterial species tree discordance test, it doubled in some settings (Supplementary Figures S17 and S18). The overall accuracy was also significantly increased, making SonicParanoid2 the most accurate based on three classification methods provided by the QfO benchmark service (Figure 2d and Supplementary Figures S5-8). More importantly, such boost in accuracy is higher when SonicParanoid2 is executed at lower sensitivities, allowing for accurate predictions with shorter runtimes (Supplementary Table S8).

The recovery of eukaryotic uridine–cytidine kinases are examples of orthologous relationships recovered using domain-based orthology inference. Previous studies showed that the fusion of prokaryotic phosphoribulokinase/uridine kinase and uracil phosphoribosyltransferase domains (P0A8F4 and P0A8F0) resulted in the emergence of this enzyme in human [20] (UniProt ID Q9NWZ5). Whereas the graph-based method of SonicParanoid2 identified P0A8F4 only as an ortholog of Q9NWZ5, the domain-based method identified P0A8F0 as well. Additionally, the domain-based method predicted P0A8F0 to be an ortholog of other 17 eukaryotic uridine– cytidine kinases in 12 eukaryotic species in the QfO dataset (Supplementary Table S9).

## Conclusion

Fast, accurate, and comprehensive identification of orthologous genes is becoming increasingly important in genomic and evolutionary studies owing to the steady growth of publicly available genomic data [14]. Nevertheless, existing *de novo* orthology inference tools require long execution times because of the necessary all-versus-all alignments and cannot effectively detect orthologs of proteins with complex domain architectures or those that have undergone gene fusion or fission events [20,21].

SonicParanoid2 solves the two aforementioned problems using ML and is faster and more accurate than existing tools. The novel graph-based algorithm almost halves the execution time by avoiding unnecessary homology searches and does not degrade the accuracy. The domain-aware algorithm increased the number of predicted orthologs by one-third and significantly increased the accuracy with minimal costs for the total execution time. Evaluation based on standardized benchmarks showed that SonicParanoid2 is the most accurate amongst other 14 well-established methods.

Considering the way SonicParanoid2 uses language models to infer orthologs, and their recent applications in bioinformatics (e.g., ProtVec [40], SeqVec [41] and ESMFold [27]), we expect accurate alignment-free orthology inference tools to be realized in the near future, which will result in significant advancements in comparative and evolutionary biology.

## Material and Methods

### Test datasets, accuracy benchmark and ranking

The execution times of SonicParanoid2, Broccoli, and OrthoFinder were evaluated using the 2020 version of a benchmark dataset provided by the QfO consortium [36]. This dataset contains 78 proteomes (Supplementary Figure S1a). Because it includes both closely and distantly related species, it provides a good example on the manner by which evolutionary relatedness affects the AdaBoost-enhanced graph-based algorithm in SonicParanoid2 (Supplementary Figure S12).

The scalability of SonicParanoid2 was evaluated on a dataset comprising 2,000 microbial MAGs from terrestrial and aquatic environments (Supplementary Figure S1b), which were randomly selected from a public catalogue of 52,515 MAGs [42]. The MAGs in the dataset satisfied or exceeded the medium quality level of the minimum information about metagenome-assembled genome (MIMAG) standard [43] and contained 5.1 million proteins. The final dataset contained 1,786 pairs of MAGs (involving 931 MAGs) with an average amino acid similarity (AAI) exceeding 99%, suggesting that for approximately a quarter of the MAGs there is another MAG which is almost identical. Although we processed these MAGs, we recommend removing highly similar proteomes from the input dataset to avoid bias in the orthology inference.

To evaluate the accuracy of SonicParanoid2, the orthologs predicted for the QfO dataset were uploaded to a benchmark service provided by the QfO. The benchmark provides 12 tests [36]: two are based on reference gene trees, two use functional information, one is based on manually curated sets of orthologous relationships among mammals, and seven assess the validity of predictions in terms of the accuracy of the species trees that can be reconstructed from them. In most of the tests, small subsets of the uploaded predictions were used, except for the LUCA generalized species tree discordance test, which evaluated all predictions.

The accuracy of SonicParanoid2 was compared with those of other 14 methods, the results of which are publicly available on the benchmark web page (as of February 2023). Benchmark results for Broccoli are not available; furthermore, because only the developers of each tool can render the benchmark results publicly available on OpenEbench [44], the accuracy of Broccoli is not discussed herein.

The ranking of the 15 methods (Figure 2d and Supplementary Figures S5-8) were done based on the three classification methods (diagonal quartiles, K-means clustering, square quartiles) offered by the QfO benchmark services. Each participant was ranked counting how many times it was placed into groups 1 to 4, where group 1 contained methods with the highest performance. The ranking shown in Figure 2d and Supplementary Figure S5 was obtained aggregating ranks from the 3 classification methods. It should be noted that the square quartiles classification rewards participants with higher precision, which resulted in rankings significantly different from those obtained from the other two classification methods.

### Software, databases, and settings

To perform a comparison of the execution times on the QfO 2020 and the MAG datasets, we used SonicParanoid 2.0.0 (commit 6698944d), Broccoli 1.2 (commit 032064c), and OrthoFinder 2.5.4 (commit 93e0e20). For the results regarding only graph-based orthology inference, SonicParanoid2 was executed with the option “skip-arch-orthologs” to omit domain-based predictions and sensitivity settings, as shown in Supplementary Table S1. OrthoFinder was executed with additional parameters, i.e., “msa” and “ot”. The former instructs OrthoFinder to use MAFFT [45] (v7.490) to compute multiple sequence alignments, whereas the latter terminates execution after the creation of HOGs [46], thus avoiding the computation of the final species tree. Broccoli was executed at the default settings.

To evaluate the AdaBoost-enhanced graph-based method, SonciParanoid2 was executed using the settings described in Supplementary Table S3, which were used in older versions of SonicParanoid (up to v1.3.8).

OrthoFinder and Broccoli were implemented in the Python3 programming language, whereas SonicParanoid2 was implemented in Python3 and Cython [47] (v3.0.0a10). All tests were performed using Python 3.8.10. The execution times of SonicParanoid2, Broccoli, and OrthoFinder were computed using the *Perf_counter* function in Python.

The alignment tools used in this study were MMseqs2 [15] (13-45111), Diamond [16] 2.0.12, and BLAST [17] 2.12.0. Supplementary Table S3 shows the mapping of the sensitivity of the alignment tools to the sensitivity settings of the tested orthology inference tools.

The average AAIs of proteome pairs in the QfO and MAGs datasets were computed using CompareM 0.1.2 (https://github.com/dparks1134/CompareM).

The profile database for Pfam [39] (version 35) was retrieved using the command *databases* in MMseqs2 with “Pfam-A.seed*”* as a parameter and indexed using the *createindex* command with parameters “-k = 5” and “-s = 7”.

The Doc2Vec model was trained using the Gensim [48] (ver. 4.2) library for Python3.

### Graph-based orthology inference

The workflow of graph-based orthology inference (Figure 1a) for *N* input proteomes is shown in Supplementary Figure S11. The first step involves predicting the *N(N-1)/2* fastest inter-proteome alignments using the AdaBoost binary classifier. Next, local alignments of the proteome pairs predicted to have the shortest execution time are performed using all input proteomes. For each of these alignments, let us assume A-B, the hits are processed to create two essential subsets (*ess(A)* and *ess(B)*) of the original proteomes using the conditions shown in Equation 2. These subsets are used for the remaining *N(N-1)/2* inter-proteome alignments. *N* intra-proteome alignments are performed in parallel using complete proteome sets.

Once all local alignments are completed, their outputs are processed, and the graph-based ortholog relations are inferred using Equation 3 and clustered as in the original SonicParanoid [30].

### AdaBoost training and optimization

The AdaBoost [32] binary classifier was built using 62,250 training samples containing properties extracted from 250 reference proteomes (Supplementary Figure S19) and labeled using the execution times of inter-proteome alignments performed using MMseqs2 at the highest sensitivity. Given *N* input proteomes, for each of the *N(N-1)* inter-proteome alignments, we created a training sample with properties such as protein count, proteome size, and average protein length. Additionally, we included the differences in proteome size and protein count, expressed in folds, by assuming that the query proteome (leftmost in the pair) was smaller and had fewer proteins than the target proteome. Each sample was labeled based on the execution times of the two inter-proteome alignments of the proteome pair (Supplementary Table S2). As an example, for a sample representing A-B, if the execution time for its alignment is shorter than that of the other alignment (B-A), then its label is set to 1 (faster); otherwise, it is set to 0 (slower).

We selected the features in Supplementary Table S2 because, for two proteomes (A and B) with substantially different compositions, the execution times for the inter-proteome alignments can differ significantly. More specifically, if proteome A contains much fewer sequences (or amino acid bases) than proteome B, then the time required to align A-B may be much shorter than that to align B-A. Furthermore, because the difference in execution time was directly proportional to the difference in the composition of the proteomes, regardless of the alignment tool used (Supplementary Figures S9 and S10); this information was used as the training feature of the AdaBoost classifier. Finally, because this information is computed and used at each execution by SonicParanoid2, regardless of the binary classifier, the creation of samples to predict the fastest alignments has virtually no overhead.

Based on the accuracies of the model on Diamond and MMseqs2 (Supplementary Table S4) as well as considering the estimated execution time for labeling the training samples using BLAST (which is likely more than two months), we decided to use only the model trained on samples labeled using MMseqs2.

The training, validation, and optimization of the model were conducted using AdaBoost libraries in Scikit-learn [49]. Hyperparameter optimization was performed using a grid search, where the best performing model achieved a mean test accuracy of 97.90 in a 10-fold cross validation. For the independent QfO test dataset, the maximum accuracy achieved by the model was 96.70% (Supplementary Table S4).

### Construction of domain architectures and training corpus for Doc2Vec

Examples of architecture estimation and document creation for a single protein are depicted in Supplementary Figure S20. Hereinafter, we will use the terms “document” and “architecture” interchangeably, and similarly for “word” and “annotated/unannotated domains”. The first step in estimating the architectures involves searching for input proteins in a Pfam profile database using MMseqs2. The hits are filtered to obtain a bit score and target coverage of at least 30 and 75%, respectively. For queries with multiple domains, the architectures are composed of non-overlapping domains and inter-regions for which no annotation was identified (uncovered regions). The elements of the architectures are converted into words, and the architectures into phrases (documents). In the documents, annotated domains are represented by their Pfam annotations, and uncovered regions (longer than four amino acid) by their lengths (Supplementary Figure S20). A document constitutes the training corpus if it has a protein coverage (the proportion of the protein sequence annotated with domains) of at least 70% and is not repeated in the corpus.

### Doc2Vec model training

Doc2Vec (also known as Paragraph2Vec) is an extension of Word2Vec [50] that represents a document as a numerical vector [33]. It uses Word2Vec to generate embeddings of single words; therefore, the embeddings can be learned through a continuous bag-of-words or skip-gram algorithm. In Doc2Vec, an algorithm that is similar to skip-gram is known as the “distributed bag-of-words of the paragraph vector”, and is used to train the Doc2Vec models in SonicParanoid2. The context window value for the skip-gram was set to two, the number of dimensions (vector size) was set to 100, and training was performed for 200 epochs. Although these parameter settings are typically used in studies pertaining to NLP [51], we set the minimum word frequency to one. This parameter controls the minimum number of times a certain word must appear in the corpus in order to be used for training. Because NLP models are typically trained on billions of documents [22], this parameter is often set to values equal to or higher than five to reduce the training time and memory usage. Nevertheless, in our case, the training sets were relatively small, where 134,520 and 567,119 documents were obtained for the QfO and MAG datasets, respectively. Hence, we set the minimum word frequency to one, allowing the neural networks of the Doc2Vec model to be trained using all documents in the corpus.

### Architecture prefiltering and creation of domain-based ortholog clusters

Given proteomes A and B with *l* and *m* architectures, respectively, *l* x *m* pairs of architectures exist. Accordingly, computing the cosine similarities for all the possible pairs of architectures for the *N(N-1)/2* combinations of input proteomes will be extremely computationally demanding. To reduce the execution time, SonicParanoid2 performs prefiltering, in which pairs of architectures that are unlikely to be orthologous are rejected before their cosine similarities are computed. Filtering is performed by comparing features of the architectures, including the protein length, protein coverage, and number of annotated domains (Supplementary Figure S20c).

A pair of architectures is rejected if one of the following conditions applies:

1. The protein length of one architecture exceed 3X the length of the other.
2. Their protein coverages differ by more than 25%.
3. One of the two architectures has more than double the domains of the other.
4. The two architectures have no mutual domains.

After prefiltering, the cosine similarities between the remaining pairs of architectures are computed, and cosine similarity values of at least 0.5 populate a matrix M of dimension *l* x *m*. The domain-based algorithm generates clusters of orthologs by selecting the maximum cosine similarities in M for proteins from A (row-wise) and B (column-wise) proteomes (Supplementary Figure S21).

### Merging of pairwise orthologs and inference of OGs

The pipelines depicted in Figures 1a and 1b infer pairwise orthologous relationships using the graph- and domain-based algorithms, respectively. SonicParanoid2 combines these orthologs to generate ortholog graphs, which are subsequently used to infer OGs (Figure 1c). Because we regarded graph-based predictions as more accurate, we integrated domain-based predictions into them.

Let G and D be sets of ortholog clusters for proteomes *A* and *B* generated by the graph- and domain-based algorithms, respectively. Let g ∈ G and d ∈ D be clusters of pairwise ortholog relationships. When merging the two sets of predicted orthologs, we considered the following three cases:

1. d is a completely new cluster with no proteins contained in any cluster in G.
2. All proteins in d are already predicted as orthologs in other clusters in G.
3. Some proteins in d are not in any cluster in G, and some proteins in g are not in any d ∈ D.

In Case 1, the architecture of each ortholog in d must have a protein coverage of at least 75%; otherwise, it is rejected. Additionally, if even a single pair of architectures has a different number of annotated domains, cluster d is rejected completely; cluster d is added as a new cluster in G if none of the aforementioned applies.

In case 2, we prioritized graph-based predictions and use the clusters in G, which already contain the orthologs in d, without any modifications.

Case 3 is the most complicated and involves scenarios in which some proteins in d are contained in one or more clusters in G, whereas others are not. In this case, the orthologous proteins from d with coverages higher than or equal to 75% are integrated into the existing G clusters. One such example is the protein P0A8F4 (Figure 4b). For proteins A0A3B3HHD8, P0A8F4, and P0A8F0, the graph-based cluster g ∈ G contains all proteins except P0A8F0.

Cluster d contains all three proteins; therefore, P0A8F0 (with 96.2% coverage) is inserted as a new ortholog into the corresponding cluster in G.

Combined sets of pairwise orthologs are used to construct ortholog graphs, which are the input to the MCL. The output from the MCL is analyzed to extract the OGs, which is the final output of the SonicParanoid2.

### Hardware used

The results described herein were obtained using an Ubuntu 20.04.3 (Linux 5.11.0) HPC server, which was equipped with a 128 cores AMD EPYC 7742 CPU, 2 terabytes of memory, and a 3.5 terabytes solid state disk.

## Declarations

### Ethics approval and consent to participate

Not applicable

### Consent for publication

Not applicable

### Availability of data and materials

SonicParanoid2 is freely available with a GNU GPLv3 license in the Python Package Index (https://pypi.org/project/sonicparanoid) and GitLab (https://gitlab.com/salvo981/sonicparanoid2). Its documentation is available at http://iwasakilab.k.u-tokyo.ac.jp/sonicparanoid.

The benchmark results for SonicParanoid2 are publicly available at https://orthology.benchmarkservice.org/proxy/results/2020. Data and programs used to train and evaluate AdaBoost and Doc2Vec models are available at https://gitlab.com/salvo981/sonic-manuscript.

### Authors’ contributions

SC designed the program and conducted the experiments. SC and WI designed the study and wrote the manuscript.

## Acknowledgements

We are grateful to the members of the QfO consortium and all the members of the Iwasaki Laboratory for their helpful comments pertaining to this study.

---

**Figure S1:**
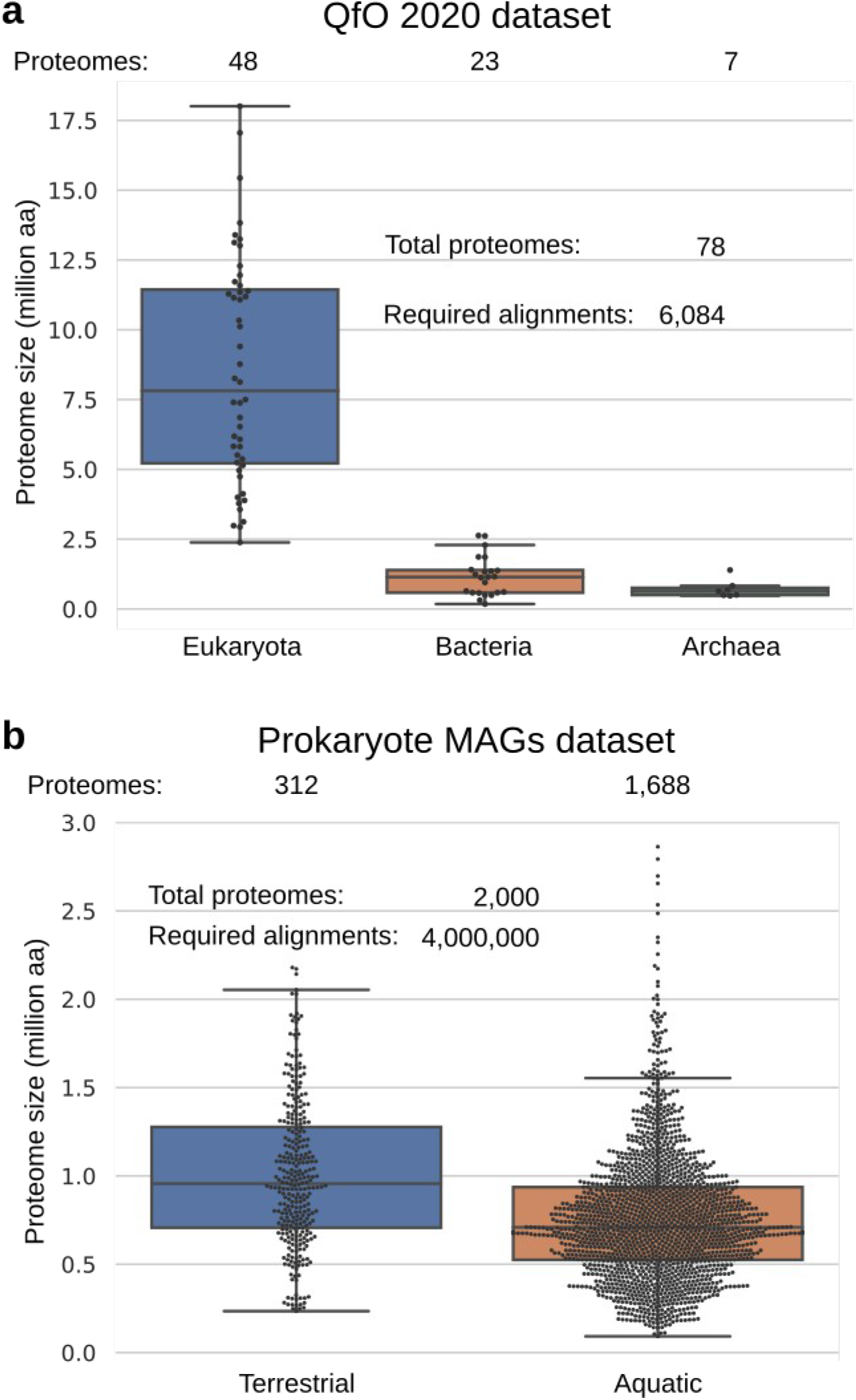
Test datasets. **a**, Version 2020 of the benchmark dataset from the QfO consortium. The dataset comprises 78 proteomes with an average size of 5.63 million AA. **b**, A set of 2,000 bacterial MAGs from terrestrial and aquatic environments randomly selected from a catalogue of 52,515 MAGs described by Nayfach *et al.* 2021.

**Figure S2:**
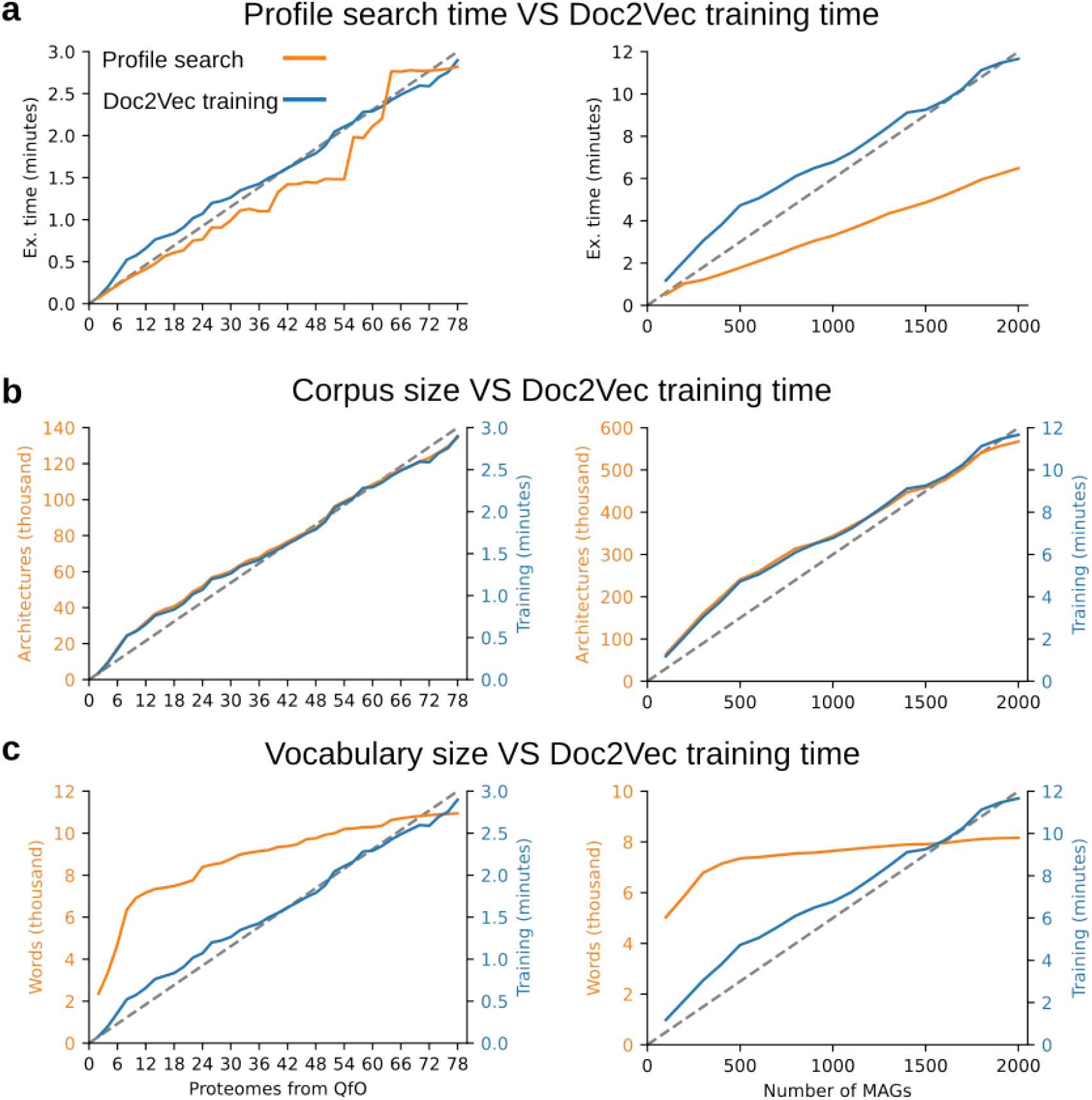
Scalability of domain-based pipeline. Plots on the right and left refer to trials with QfO and MAG datasets as input, respectively. Multiple trials were performed using the domain-based pipeline with increasing numbers of input proteomes from both datasets (x-axis). Domain-based pipeline shows high scalability with a quasi-linear execution time growth as the number of input proteomes increases. **a**, Increase in execution time for profile searches and Doc2Vec model training. Total execution time for both steps was approximately 6 min for the complete QfO dataset and approximately 18 min (only 3X longer) for the 2,000 MAGs. **b**, Time required to train the Doc2Vec models increases linearly with the corpus size. **c**, Size of vocabulary (single words in the corpus) reaches a plateau when the input proteomes have similar contents (e.g. for the MAG dataset).

**Figure S3:**
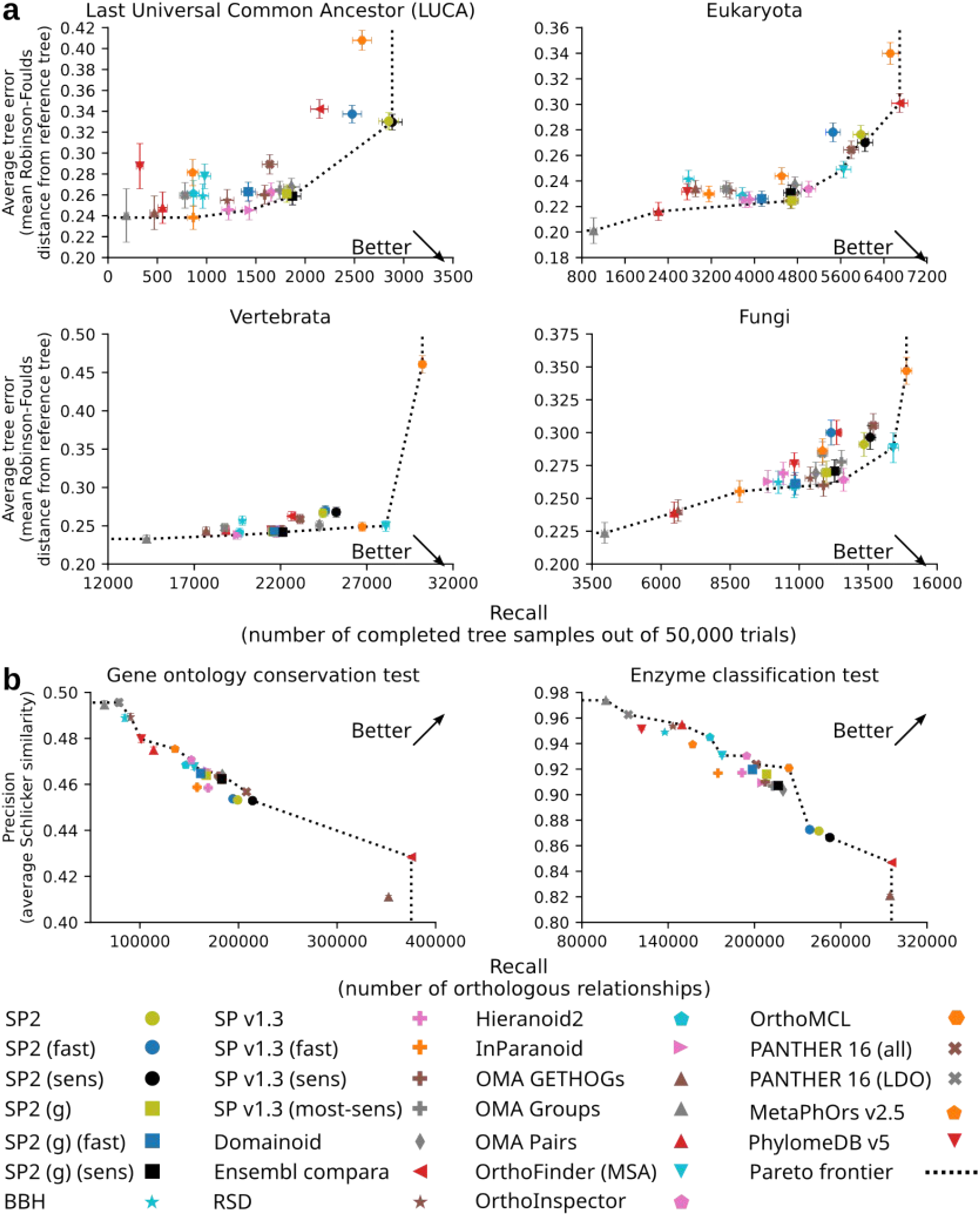
Accuracy of SonicParanoid2 with and without domain-based orthology. Methods shown by markers on the Pareto frontier demonstrate the best balance between precision and recall. Square markers represent accuracies obtained when using only the graph-based pipeline. Results of **a**, generalized species tree discordance tests and **b**, functional benchmark tests.

**Figure S4:**
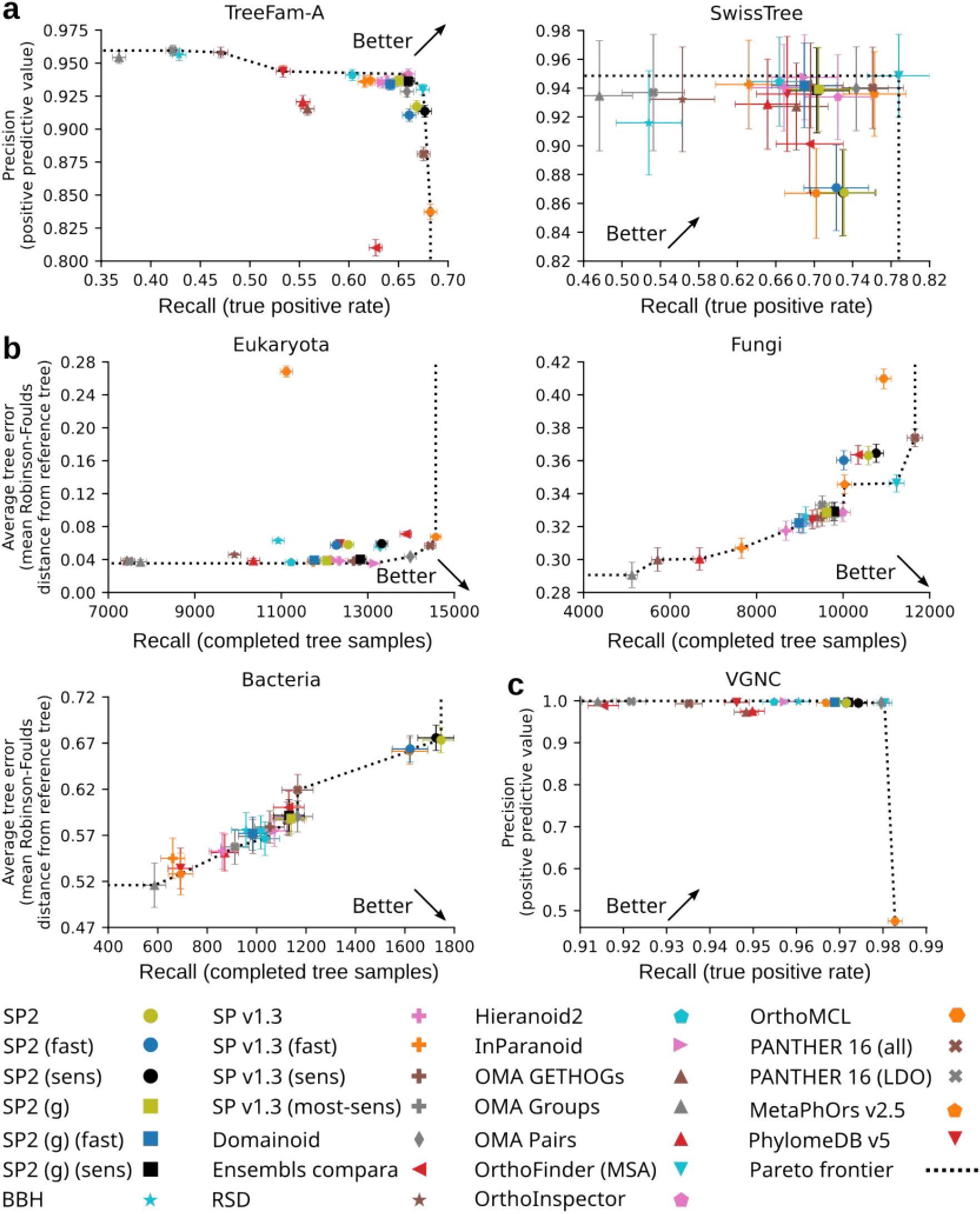
Accuracy of SonicParanoid2 with and without domain-based orthology. Methods shown by markers on Pareto frontier demonstrate the best balance between precision and recall. Square markers represent accuracies obtained when using only the graph-based pipeline. Results of **a**, reference-tree based tests, **b**, species tree discordance tests (eukaryota, fungi, bacteria), and **c**, Vertebrate Gene Nomenclature Committee (VGNC) test.

**Figure S5:**
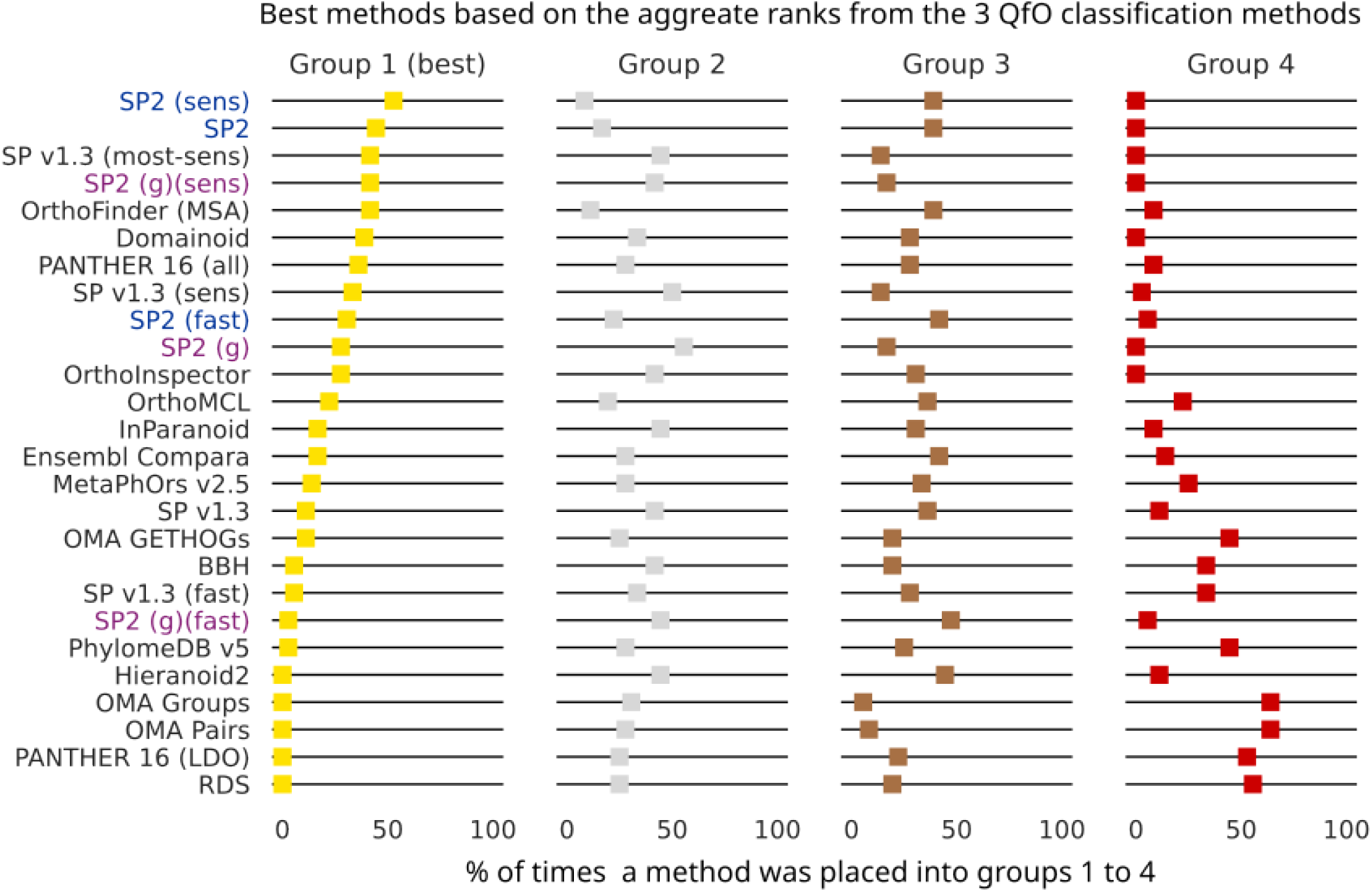
Accuracy of SonicParanoid2 and other 14 methods based on the aggregate ranking from the three classification methods of the QfO benchmark. For each test in the benchmark suite, a method was assigned to group 1 if it had very high precision and recall (e.g., close to “optimal performance” corner), and group 4 if it was close to the “suboptimal performance” corner. Based on a total of 36 rankings (12 from each classification method) participants were sorted on the percentages of tests they were assigned to each group. Blue labels represent SonicParanoid2 executions which included the domain-based orthology inference, whereas purple labels refer to runs in which only graph-based orthology was performed.

**Figure S6:**
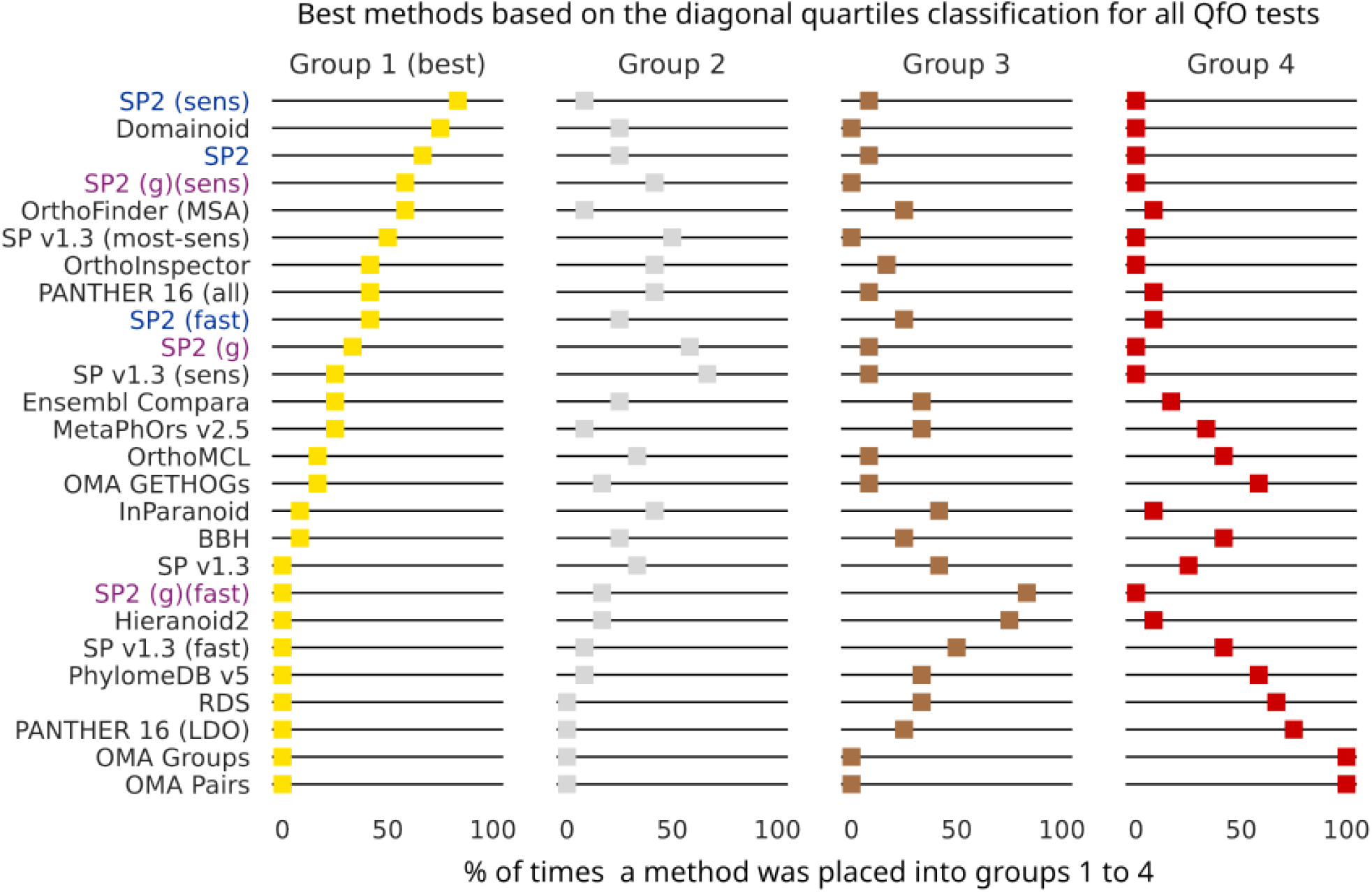
Accuracy of SonicParanoid2 and other 14 methods based on diagonal quartiles classification from the QfO benchmark. For each of the 12 tests, participants were assigned a group based on their distance from the “optimal perfomance” corner. Methods were sorted on the percentages of tests they were assigned to each group. Blue labels represent SonicParanoid2 executions which included the domain-based orthology inference, whereas purple labels refer to runs in which only graph-based orthology was performed.

**Figure S7:**
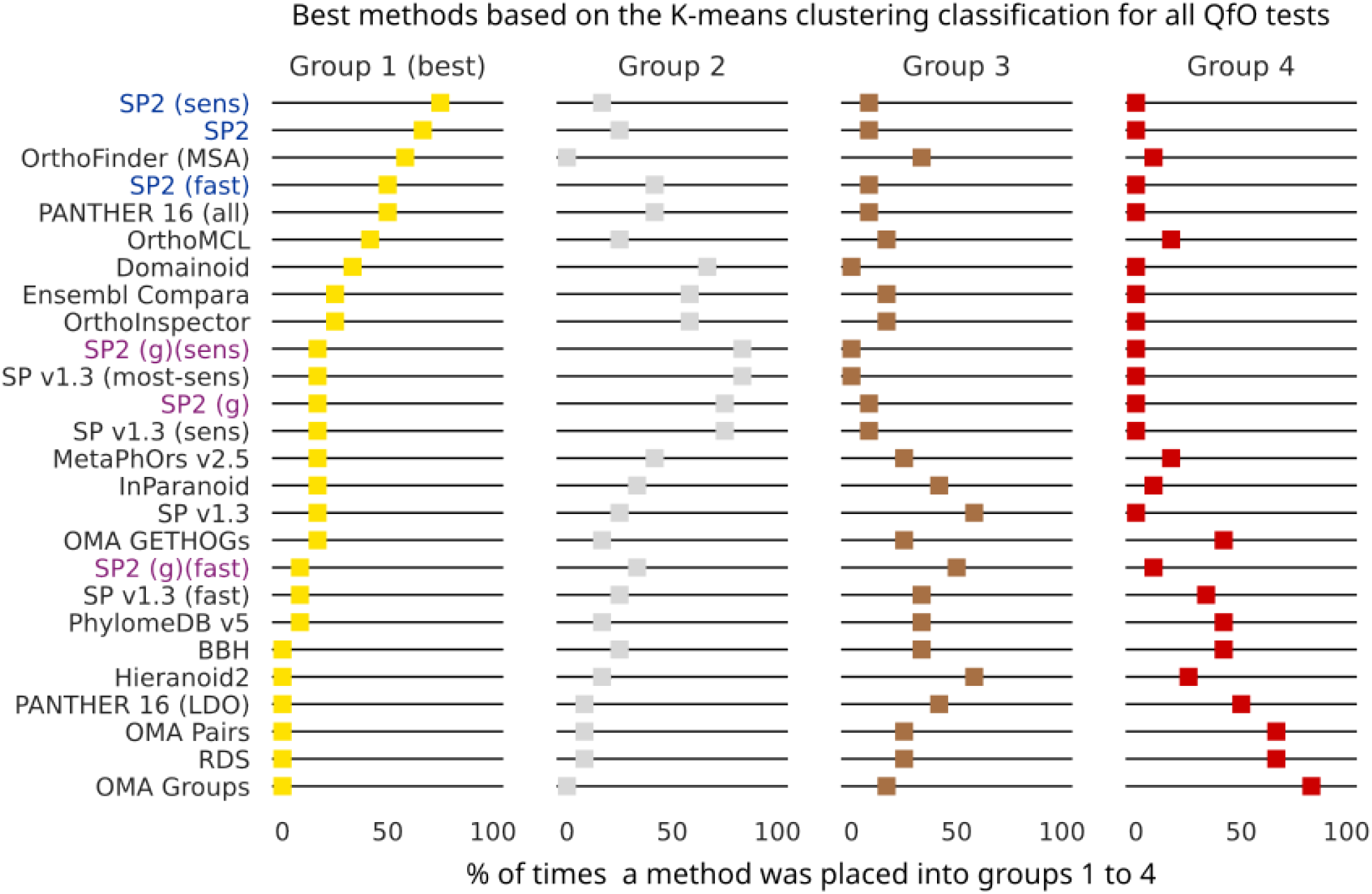
Accuracy of SonicParanoid2 and other 14 methods based on K-means clustering classification from the QfO benchmark. For each of the 12 tests, participants were separated into 4 clusters (groups) using the K-means algorithm. Methods were sorted on the percentages of tests they were assigned to each group. Blue labels represent SonicParanoid2 executions which included the domain-based orthology inference, whereas purple labels refer to runs in which only graph-based orthology was performed.

**Figure S8:**
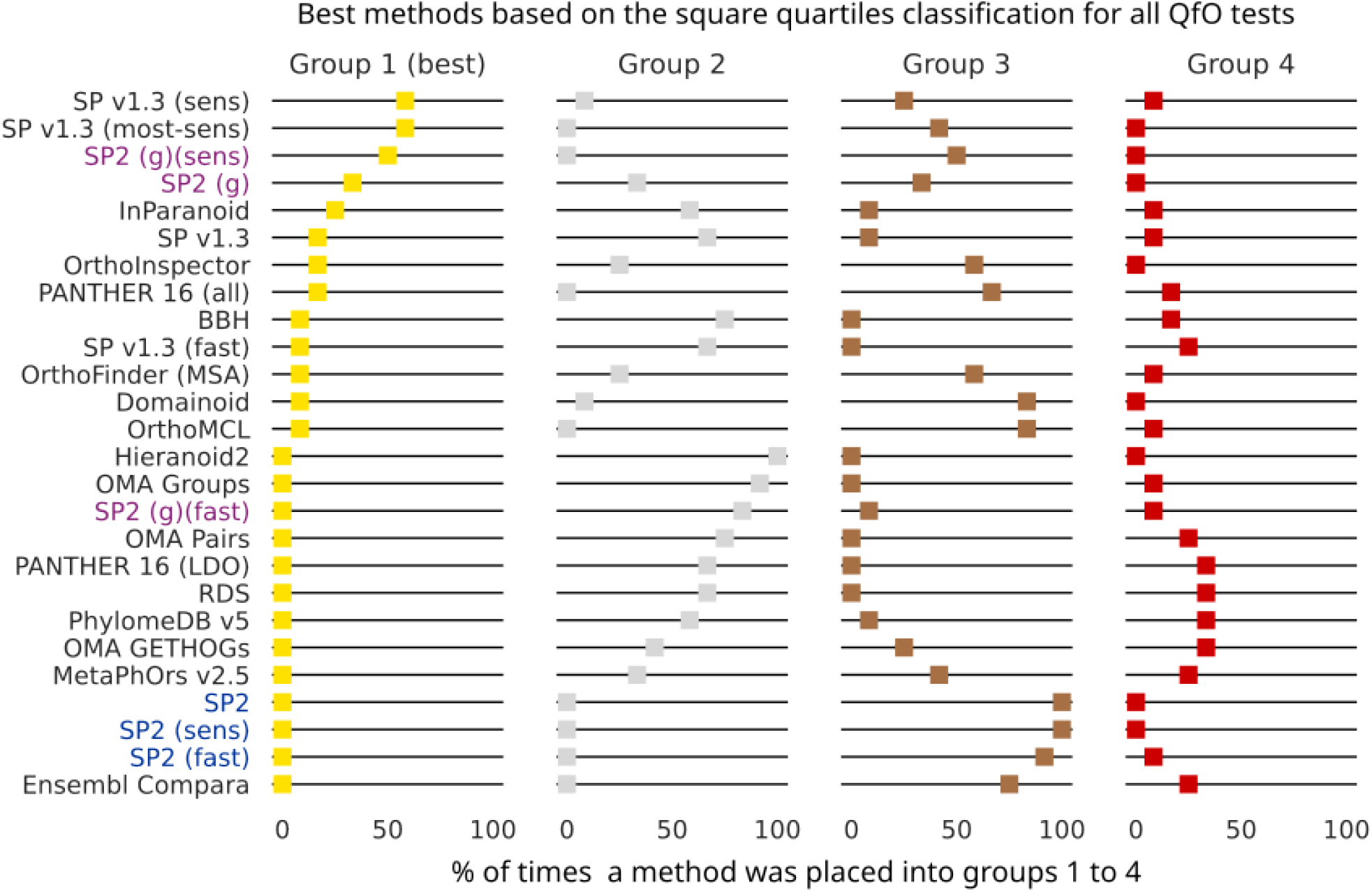
Accuracy of SonicParanoid2 and other 14 methods based on the square quartiles classification from the QfO benchmark. For each of the 12 tests, participants were assigned a group by checking if they were above or below of the first quartiles of the precision and recall. Methods were sorted on the percentages of tests they were assigned to each group. Blue labels represent SonicParanoid2 executions which included the domain-based orthology inference, whereas purple labels refer to runs in which only graph-based orthology was performed.

**Figure S9:**
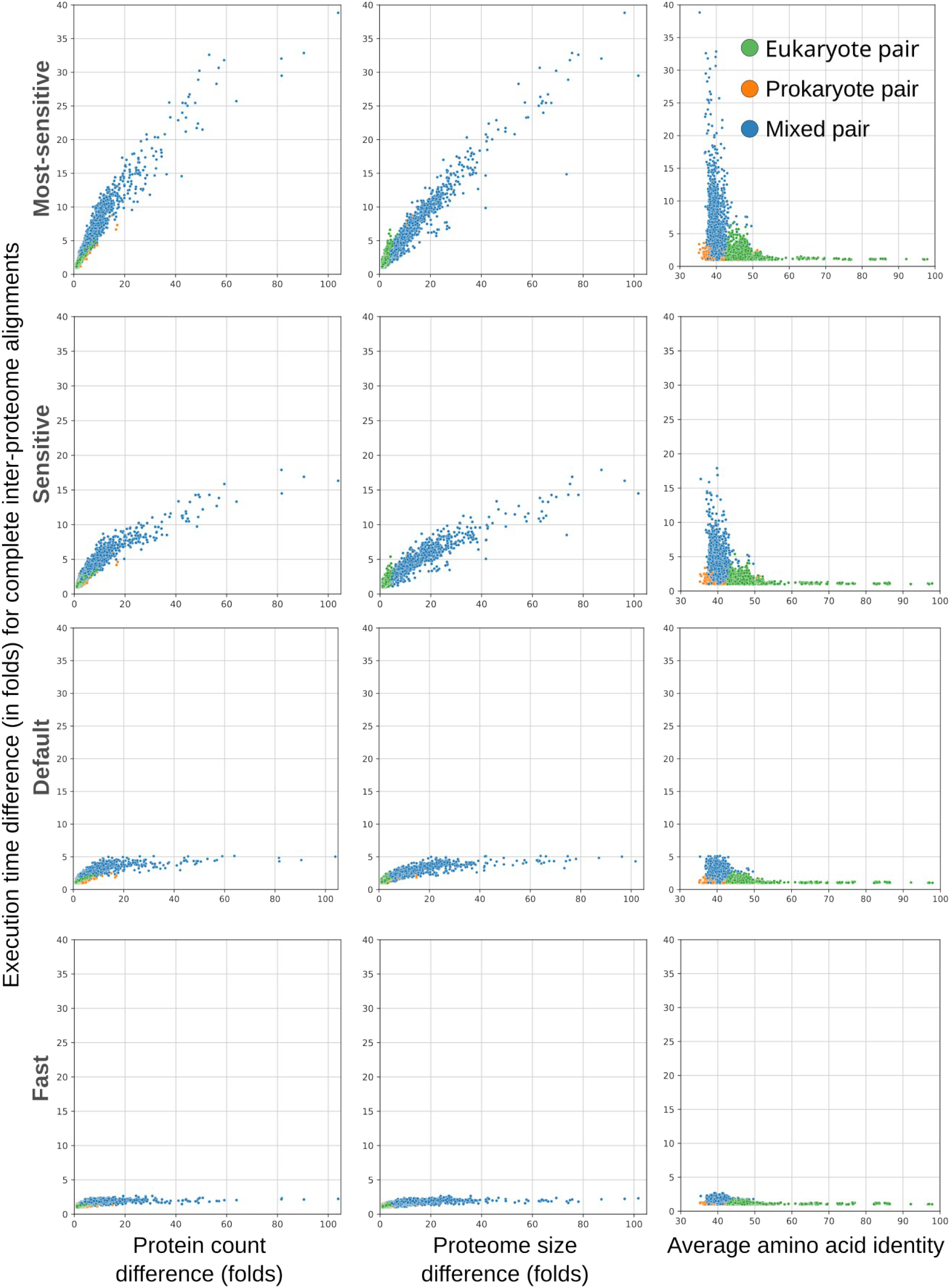
Execution time differences between complete inter-proteome alignments using MMseqs2 at multiple settings. The execution time difference for a specified pair (e.g., A-B) is computed as the absolute value of the execution times of the two inter-proteome alignments. Each dot represents one of the 3,003 possible combinations for the QfO dataset.

**Figure S10:**
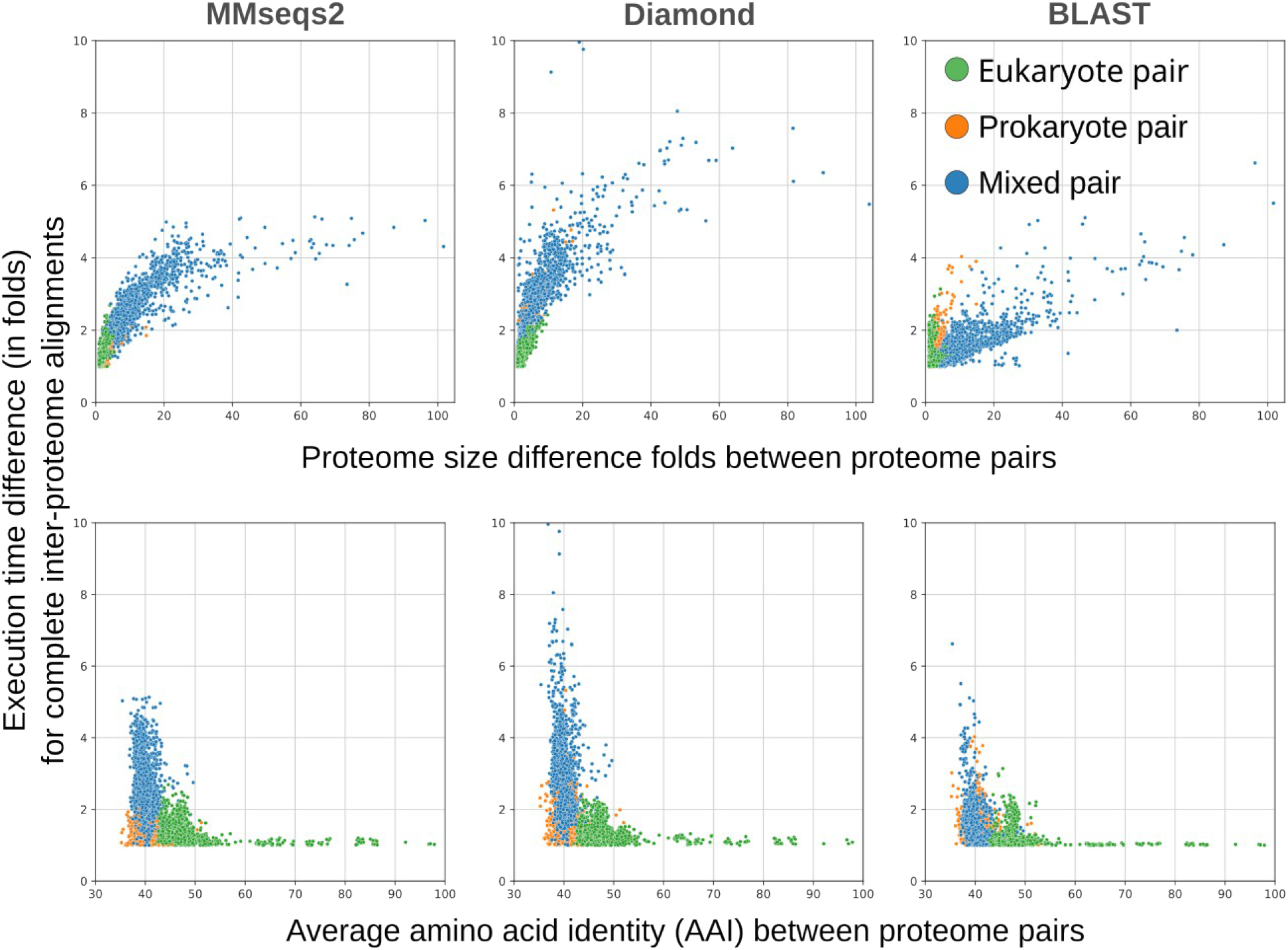
Execution time differences between complete inter-proteome alignments using multiple local alignment tools. Y-axis shows execution time difference between inter-proteome alignments for each of the 3,003 possible combinations in the QfO proteome set, using (left to right) MMseqs2, Diamond, and BLAST at the default mode (as in Supplementary Table S3). As in Figure S9, the execution time difference is proportional to the difference in proteome size (top row) and inversely proportional to AAI (bottom row).

**Figure S11:**
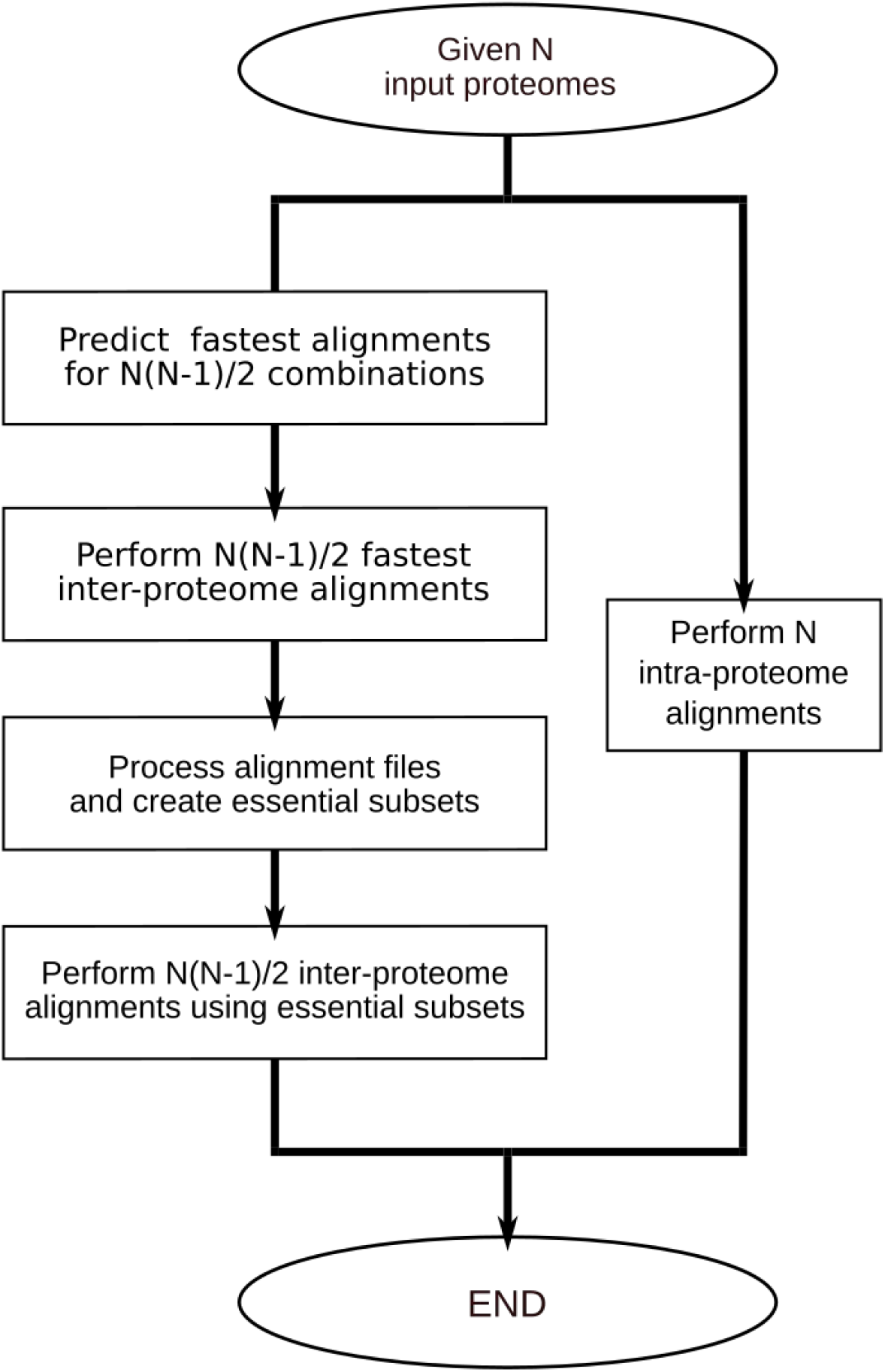
Flowchart of AdaBoost-enhanced graph-based method in SonicParanoid2. For each possible combination of *N* input proteomes, the fastest inter-proteome alignments are predicted using an AdaBoost binary classifier (left branch). After the fastest alignments are performed, they are processed and the essential protein sets are generated, as in Equation 2. The remaining inter-proteome alignments are performed on the reduced protein sets. The intra-proteome (right branch) and the predicted fastest inter-proteome alignments are performed using the complete input proteomes.

**Figure S12:**
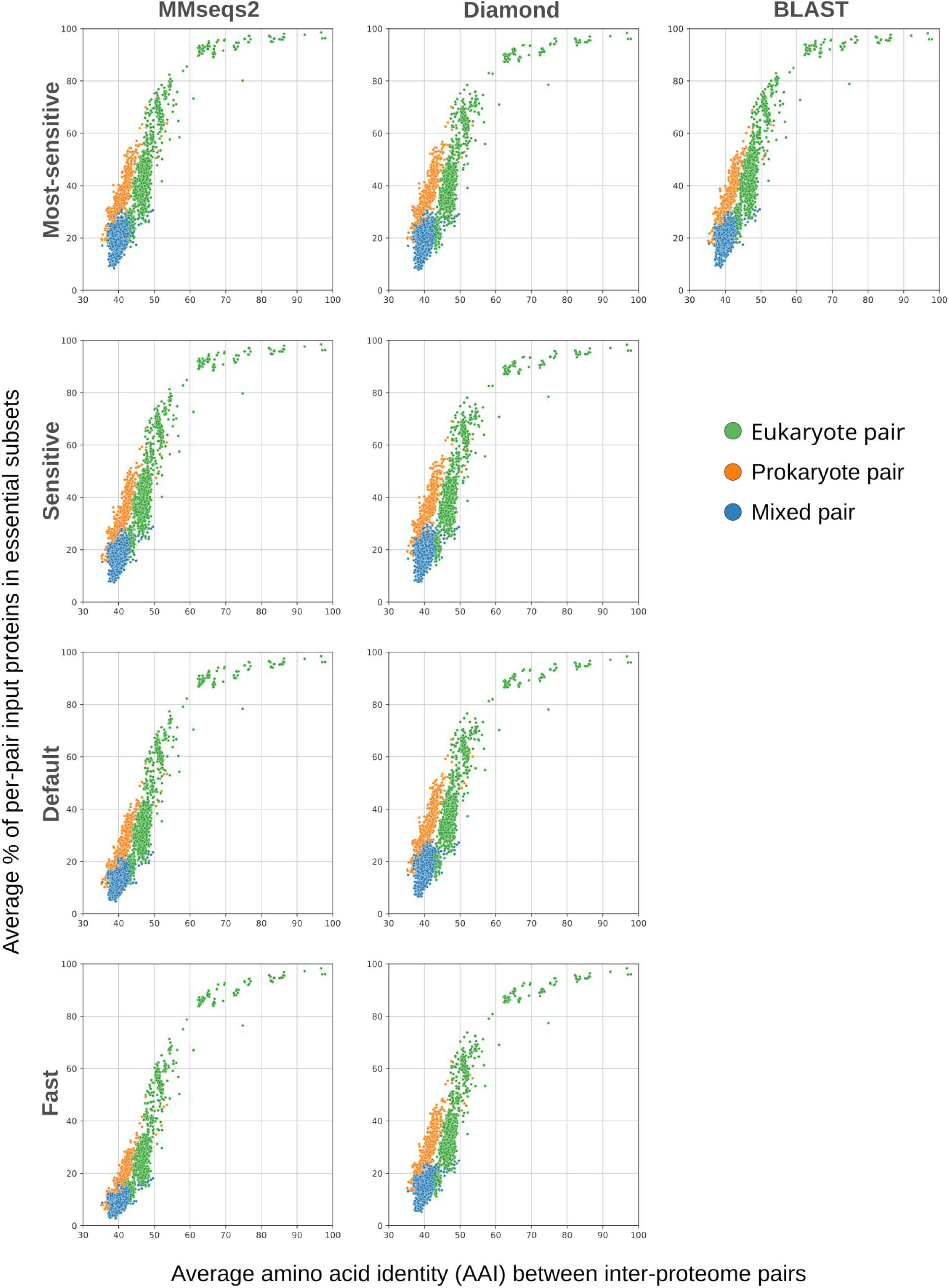
Percentage of input sequences used and evolutionary relatedness in graph-based orthology inference (QfO 2020). Y-axis shows the average percentage of original input constituting the essential subsets of each pair of proteomes. For example, if the essential sets for A and B contain 15% and 25% of the original input proteins, respectively, then 20 will be on the y-axis. The percentage of the original input proteins used is directly proportional to the evolutionary relatedness (expressed in terms of the AAI on the x-axis) and agnostic on the alignment tool used.

**Figure S13:**
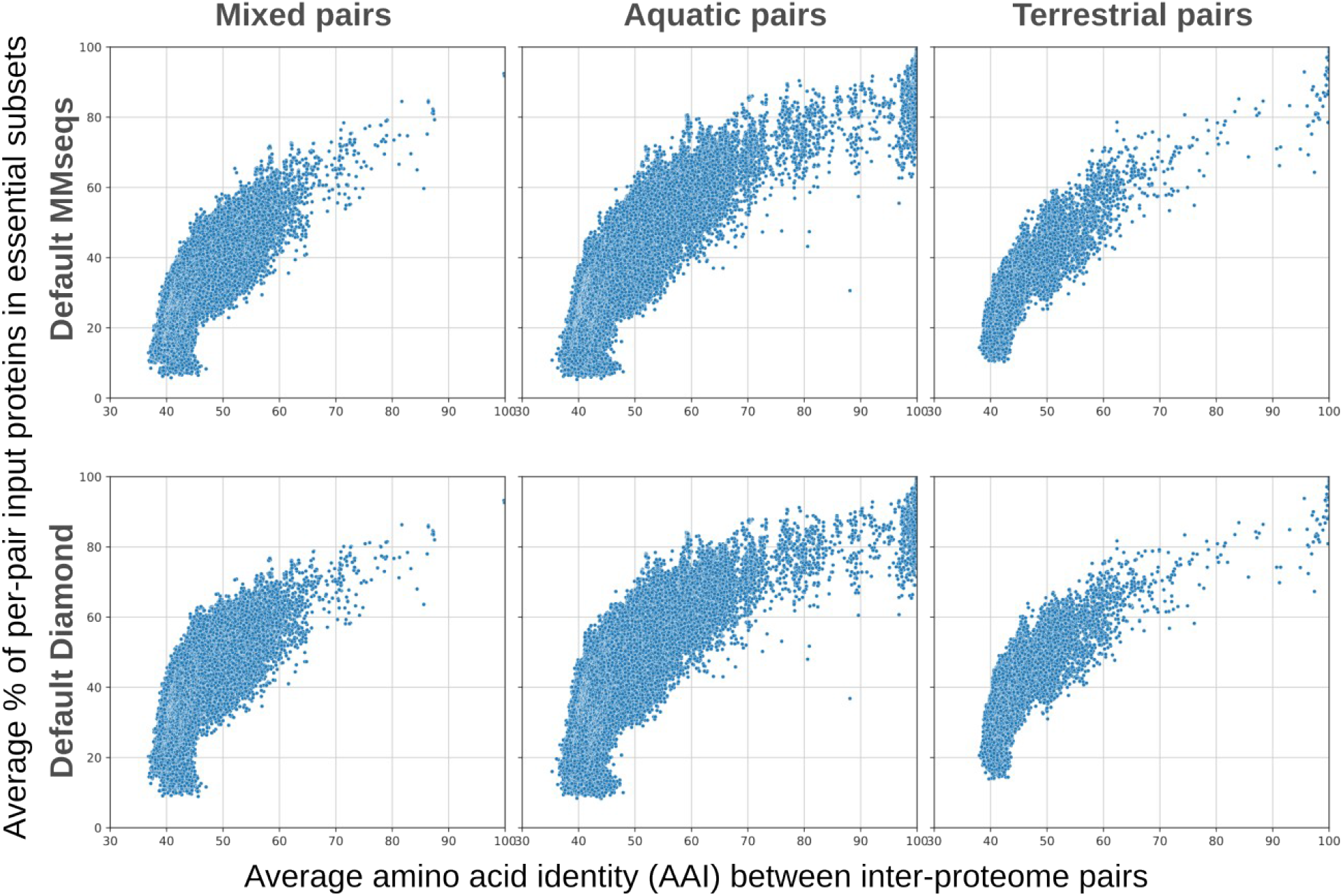
Percentage of input sequences used and evolutionary relatedness in graph-based orthology inference (2,000 bacterial MAGs). Y-axis shows average percentage of input proteins constituting the essential subsets of each inter-proteome alignment. Similarly to the observations for the QfO dataset (Supplementary Figure S12), the average percentage of input proteins used in the essential subsets is directly proportional to the evolutionary relatedness (expressed as AAI). The leftmost plots refer to pairs of MAGs, in which one MAG was obtained from aquatic samples and the other from terrestrial samples.

**Figure S14:**
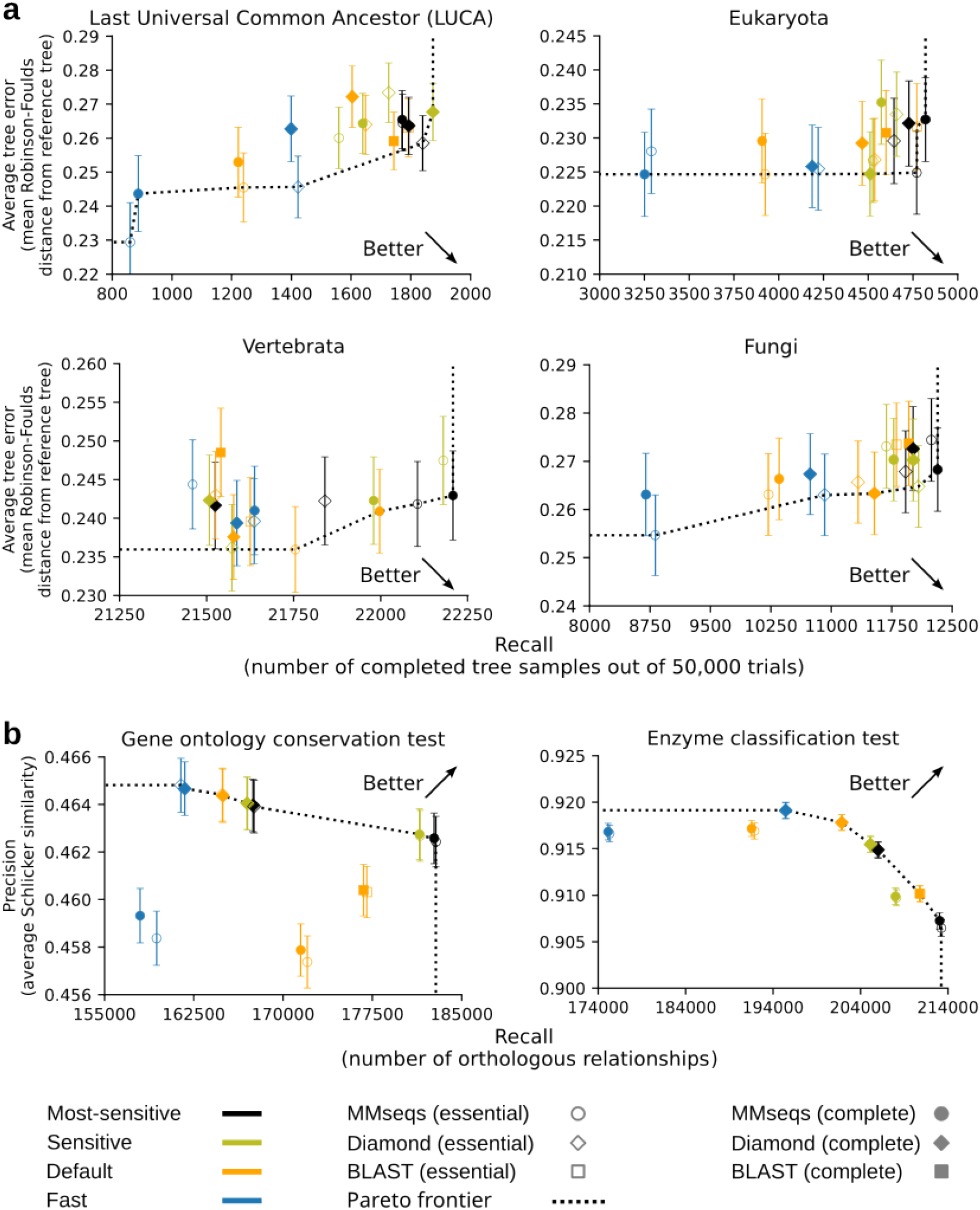
Benchmark results for predictions obtained with and without essential alignments. **a**, Accuracies of SonicParanoid2 obtained using essential alignments (empty markers) on the generalized species tree discordance tests are highly similar to those obtained using the complete alignments, regardless of the alignment tool and sensitivity settings used. **b**, The prediction accuracies on functional benchmark tests conducted using the essential and complete alignments are extremely similar and do overlap in many cases.

**Figure S15:**
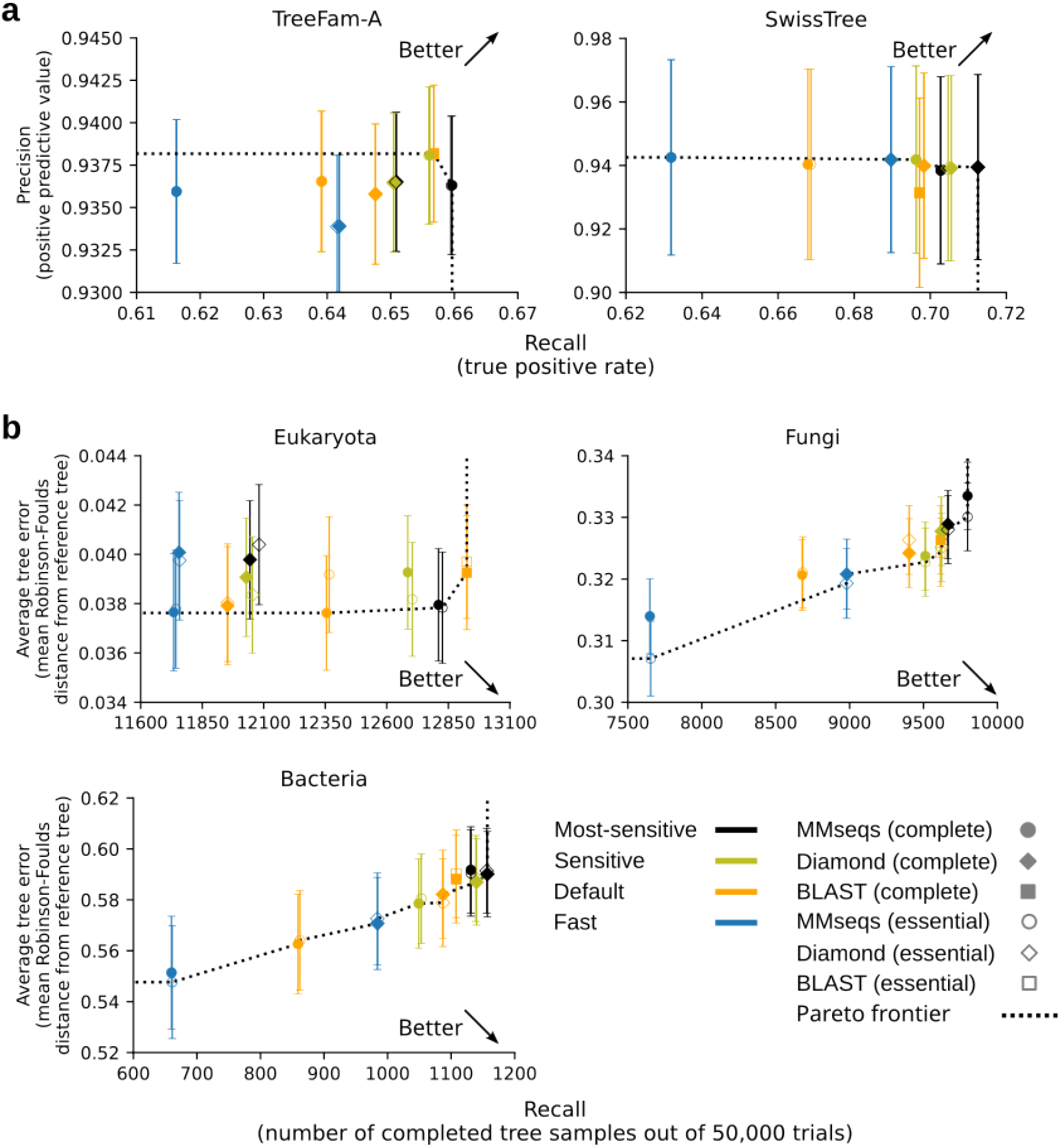
Benchmark results for predictions obtained with and without essential alignments. Empty markers represent predictions obtained using essential alignments. **a**, Prediction accuracies on gene-tree-based benchmark tests tend to overlap, which is likely due to the small sizes of the benchmark sets. Accuracies overlap on species tree discordance tests in **b** for fungi and bacteria, which use relatively small test sets. Accuracies based on eukaryote dataset appear more similar at higher sensitivity settings.

**Figure S16:**
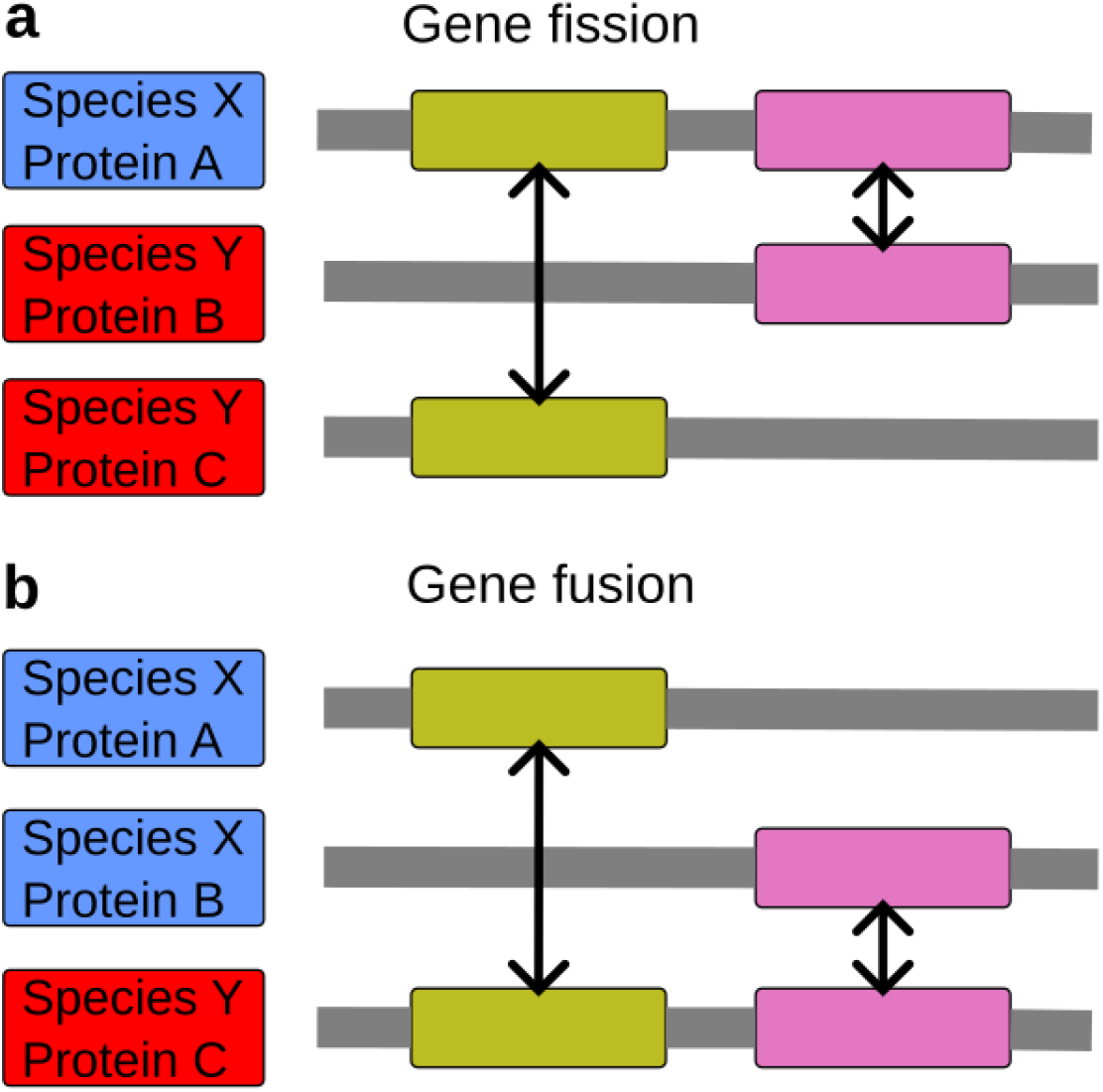
Orthologous domains typically missed by BBH. Proteins involved in fission or fusion events during evolution may contain orthologous relationships at the domain level (black arrows) that may be missed by BBH. **a**, Gene fission event in which a multi-domain protein from species X has its orthologous domains separated into two single-domain proteins in species Y. **b**, Gene-fusion event in which two domains in two different proteins in species X have their orthologs arranged into a single protein in species Y.

**Figure S17:**
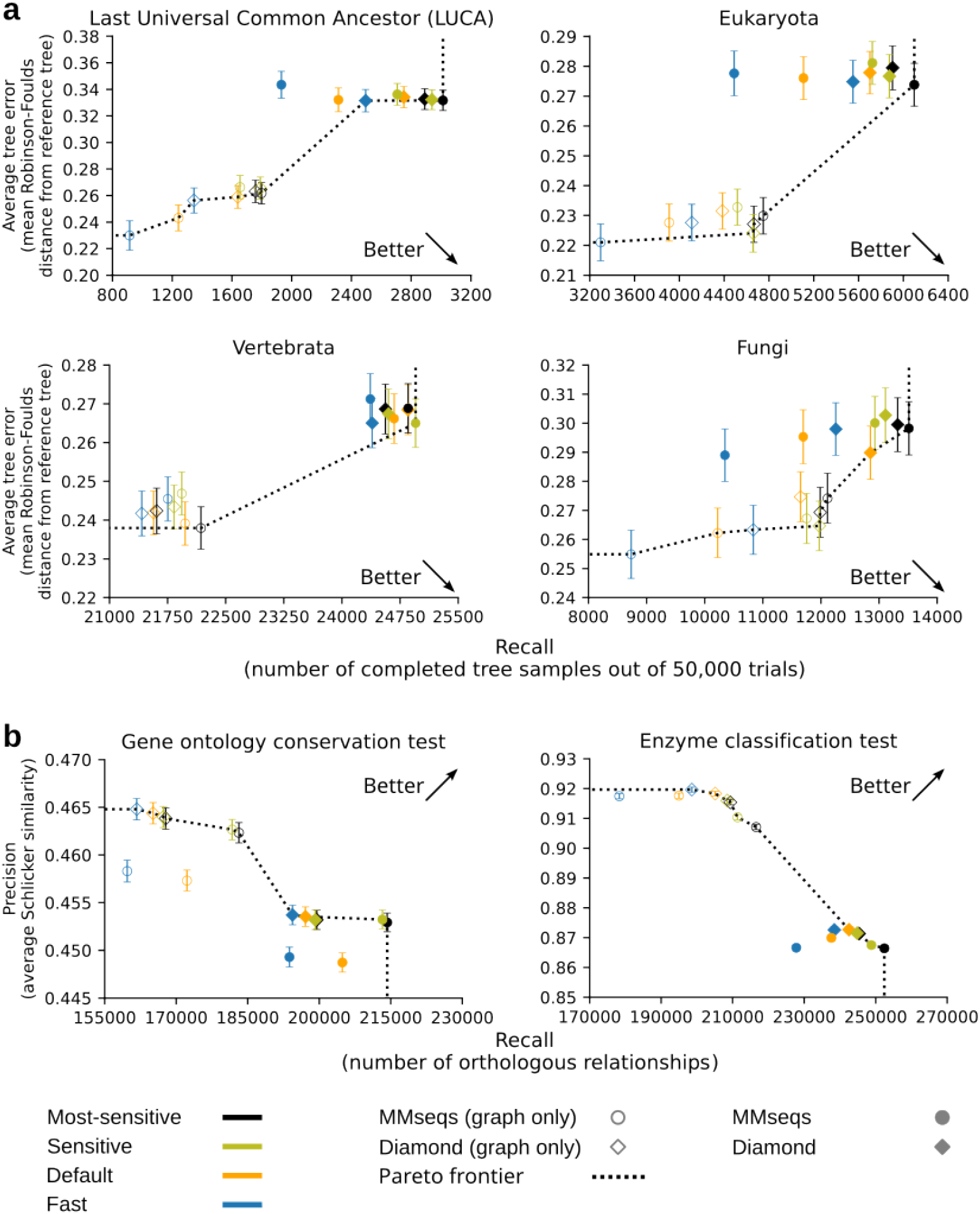
QfO benchmark results for SonicParanoid2 with and without domain-based orthology inference. Results shown by empty markers were obtained using the graph-based pipeline only (Figure 1a), whereas those shown by filled markers include orthologs predicted from the domain-based algorithm (Figure 1b). Settings described in Supplementary Table S3 were used. **a**, Accuracy results obtained from generalized species tree discordance tests. **b**, Prediction accuracies obtained from functional benchmark tests.

**Figure S18:**
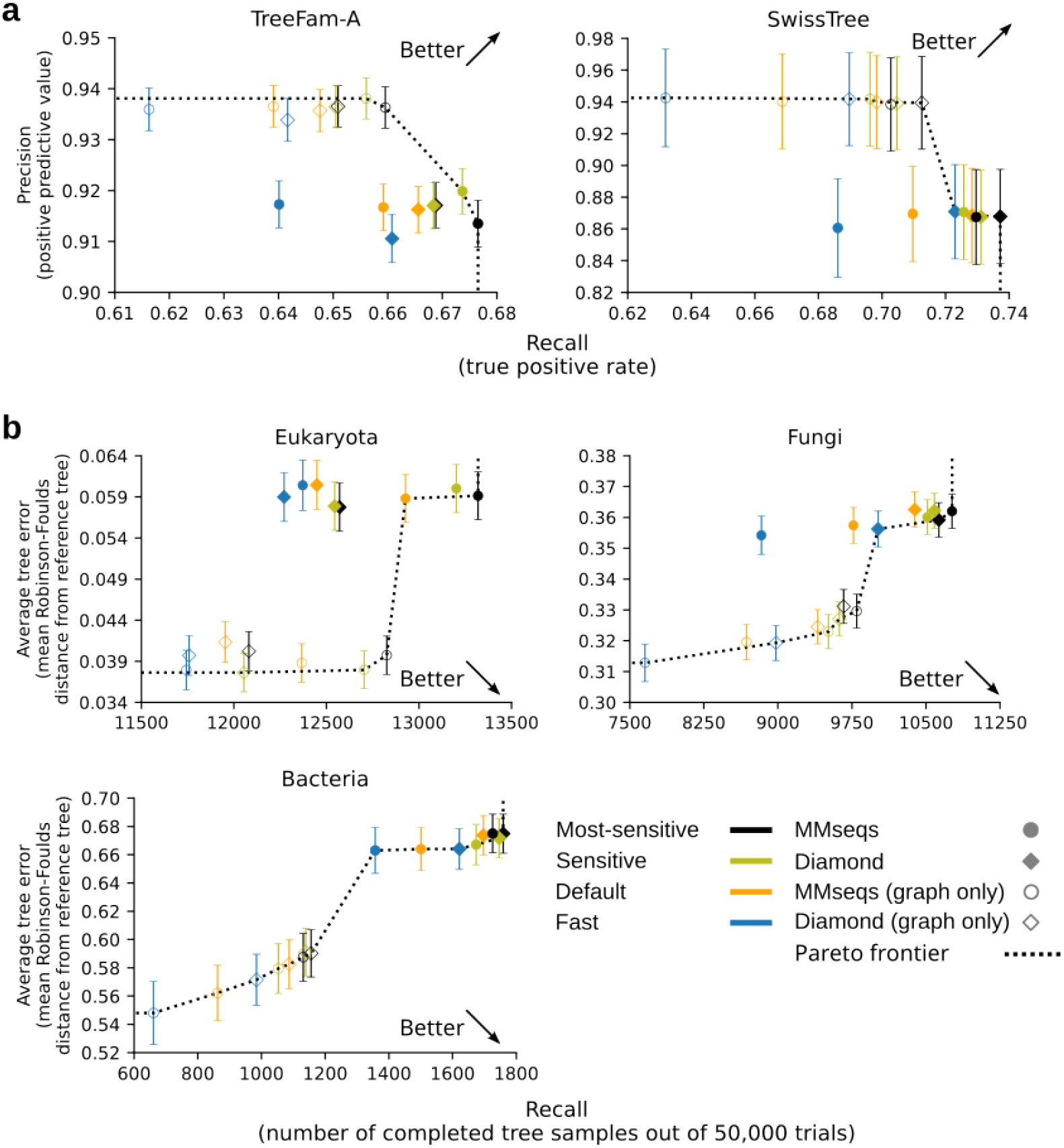
QfO benchmark results for SonicParanoid2 with and without domain-based orthology inference. Results shown by empty markers were obtained using the graph-based pipeline only (Figure 1a), whereas those shown by filled markers include orthologs predicted from the domain-based algorithm (Figure 1b). Settings described in Supplementary Table S3 were used. Results obtained from **a**, gene tree-based tests and **b**, species tree discordance tests.

**Figure S19:**
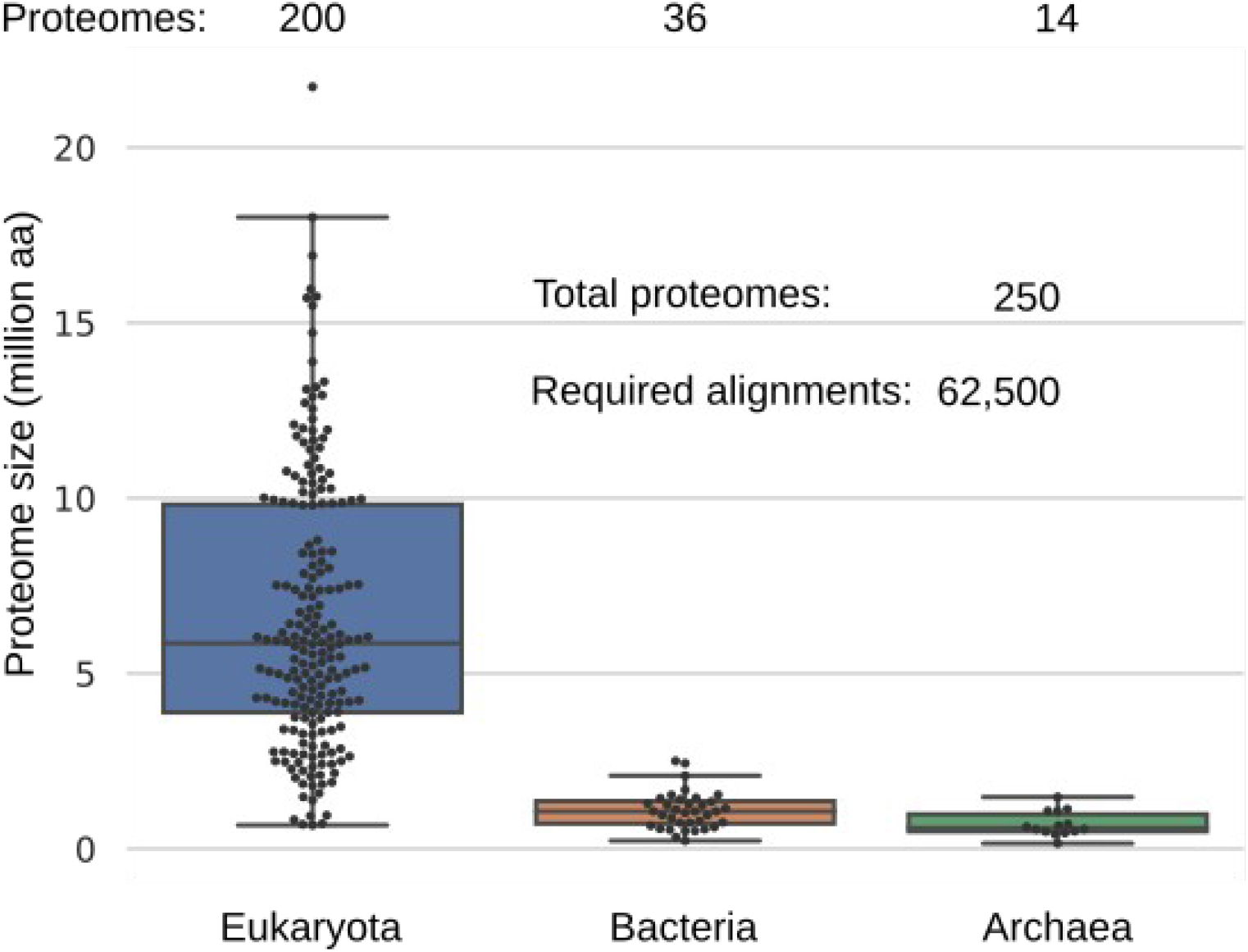
Dataset used to generate training samples and labels for binary AdaBoost classifier. Average size of the proteomes is 5.50 million AA, with *Glicine max* (21.73 million AA) and *Nanoarchaeum equitans* (0.15 million AA) being the largest and smallest, respectively.

**Figure S20:**
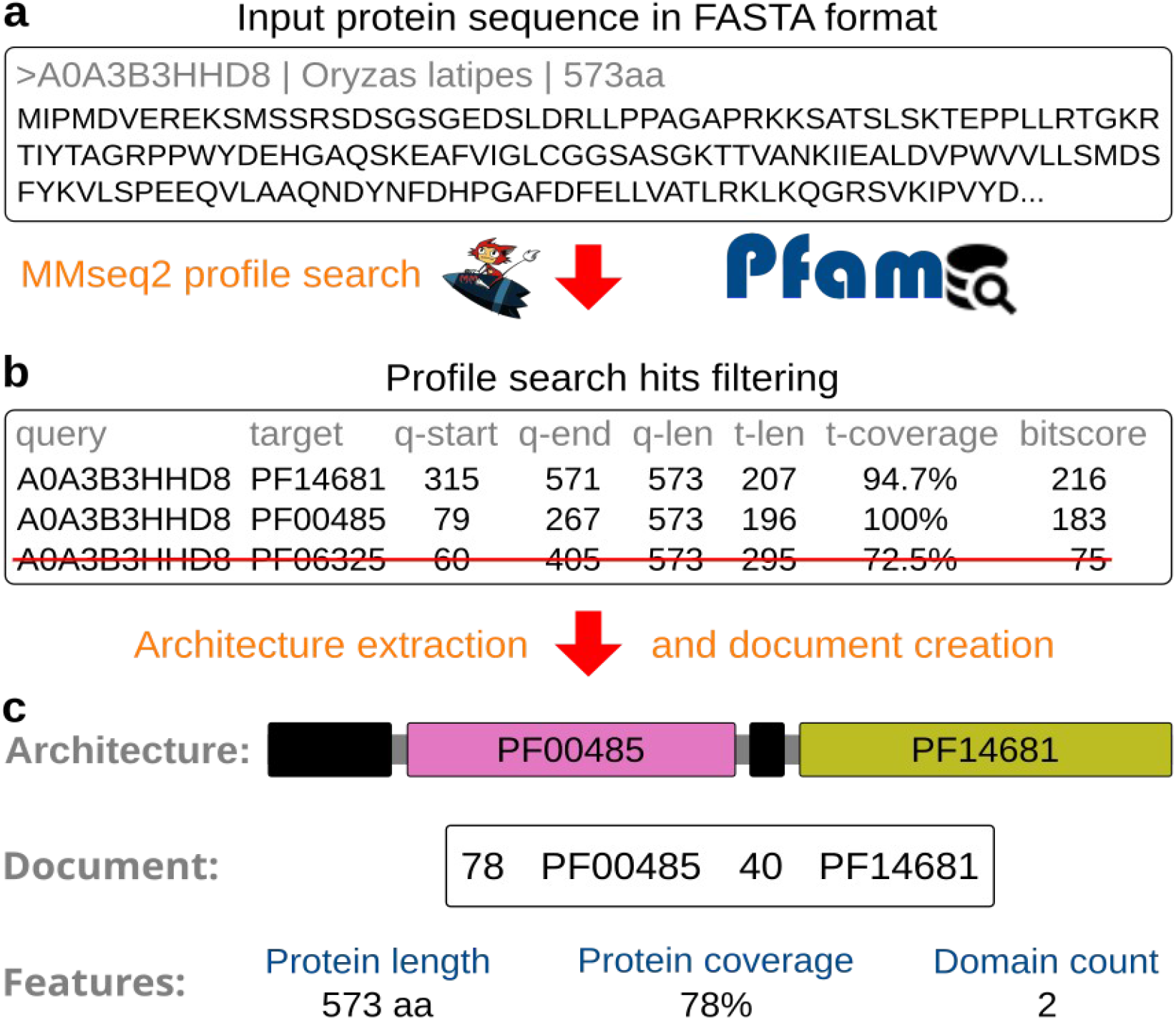
Example of architecture estimation and document creation for a single input protein. **a**, Search the protein in the Pfam profile database using MMseqs2. **b**, Only hits with a bitscore exceeding 30 and domain coverage (t-coverage) exceeding 75% are used to build the architectures. **c**, Representation of the architecture as a sequence of domains, document, and architecture features. Black blocks in the architecture represent unannotated regions.

**Figure S21:**
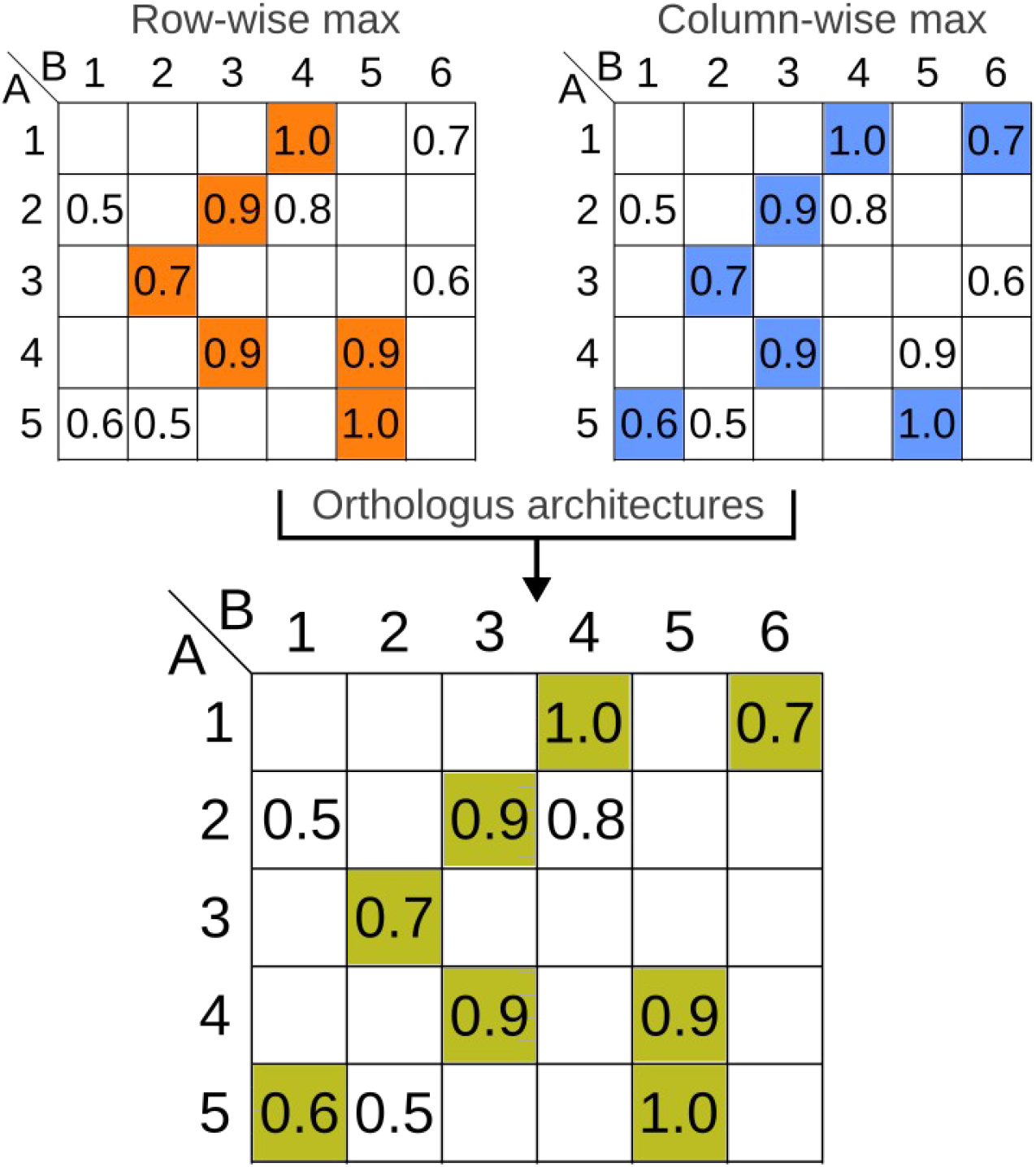
Selection of candidate orthologs in the domain-based pipeline. Maximum cosine similarities for each protein from proteomes A (top-left) and B (top-right) are selected. Proteins corresponding to entries in the final matrix (bottom) will populate the ortholog clusters generated by the domain-based pipeline.

**Table S1:** SonicParanoid2 presets. Alignment tools and settings used for the three SonicParanoid2 presets. In addition to the presets in the table, users can execute SonicParanoid2 using custom settings for the alignment tool (Diamond, MMseqs2 or BLAST) and its sensitivity.

| Mode | Alignment tool | Settings |
| --- | --- | --- |
| Default | Diamond | very-sensitive |
| Fast | Diamond | mid-sensitive |
| Sensitive | MMseqs | -s 7.5 |

**Table S2:** AdaBoost training samples. The first column shows the pairs of proteomes each sample represents (not used in the training). Columns 2–9 show training features for AdaBoost, which include information extracted from the input proteomes, such as protein count, proteome size, and average protein length. Last column shows the training labels.

| Pair | Sequence count sp1 | Sequence count sp2 | Sequence count difference folds (assuming sp1<sp2) | Proteome size sp1 | Proteome size sp2 | Proteome size difference folds (assuming sp1<sp2) | Mean protein length A | Mean protein length B | Fastest |
| --- | --- | --- | --- | --- | --- | --- | --- | --- | --- |
| A-B | 3,422 | 2,236 | -1.53 | 769,203 | 739,153 | -1.04 | 224.78 | 330.57 | 0 |
| B-A | 2,236 | 3,422 | 1.53 | 739,153 | 769,203 | 1.04 | 330.57 | 224.78 | 1 |
| A-C | 3,422 | 2,353 | -1.45 | 769,203 | 637,663 | -1.21 | 224.78 | 271.00 | 0 |
| C-A | 2,353 | 3,422 | 1.45 | 637,663 | 769,203 | 1.21 | 271.00 | 224.78 | 1 |
| A-D | 3,422 | 1,889 | -1.81 | 769,203 | 537,042 | -1.43 | 224.78 | 284.30 | 0 |
| D-A | 1,889 | 3,422 | 1.81 | 537,042 | 769,203 | 1.43 | 284.30 | 224.78 | 1 |

**Table S3:** Mapping of sensitivities of alignment tools to sensitivity settings of the tested orthology inference tools. Cells filled with “X” represent unavailable or untested settings.

| Tool | Version | Mode | MMseqs | Diamond | BLAST |
| --- | --- | --- | --- | --- | --- |
| SonicParanoid | 2.0.0 | Fast | -s 2.5 | mid-sensitive | X |
| SonicParanoid | 2.0.0 | Default | -s 4.0 | sensitive | default |
| SonicParanoid | 2.0.0 | Sensitive | -s 6.0 | very-sensitive | X |
| SonicParanoid | 2.0.0 | Most-sensitive | -s 7.5 | ultra-sensitive | X |
| Broccoli | 1.2.1 | Default | X | more-sensitive | X |
| OrthoFinder | 2.5.4 | Default | X | more-sensitive | X |

**Table S4:** Saved execution time using AdaBoost-enhanced graph-based algorithm when processing QfO 2020 dataset. Results for 18 trials of SonicParanoid2 using the graph-based pipeline are shown. Sub-tables show the results obtained using (from top to bottom) MMseqs2, Diamond, and BLAST at different sensitivities. Columns two and three show the time required to perform homology searches (in hours) using the conventional all-versus-all (complete alignments) and the novel approach (essential alignments), respectively. Accuracies for the AdaBoost classifier and the average sizes of the original input proteins used in the essential subsets are shown in columns four and five, respectively. The last two columns on the right show the amount of time saved using the novel graph-based approach (in hours and percentage, separately).

| MMseqs2 |  |  |  |  |  |  |
| --- | --- | --- | --- | --- | --- | --- |
| Mode | Complete alignments | Essential alignments | AdaBoost accuracy (%) | Avg. (%) used in essential sets | Saved hours | Saved time (%) |
| most-sensitive | 6.76 | 3.91 | 95.94 | 33.45 | 2.85 | 42.18 |
| sensitive | 2.57 | 1.65 | 94.91 | 31.22 | 0.91 | 35.60 |
| default | 0.64 | 0.47 | 94.61 | 25.50 | 0.17 | 25.84 |
| fast | 0.33 | 0.26 | 92.77 | 20.22 | 0.07 | 20.87 |
| Diamond |  |  |  |  |  |  |
| Mode | Complete alignments | Essential alignments | AdaBoost accuracy (%) | Avg. (%) used in essential sets | Saved hours | Saved time (%) |
| most-sensitive | 2.35 | 1.71 | 95.37 | 31.94 | 0.63 | 27.01 |
| sensitive | 0.62 | 0.45 | 95.20 | 31.42 | 0.17 | 27.69 |
| default | 0.46 | 0.30 | 96.10 | 29.71 | 0.16 | 34.29 |
| fast | 0.23 | 0.16 | 96.70 | 26.78 | 0.07 | 31.05 |
| BLAST |  |  |  |  |  |  |
| Mode | Complete alignments | Essential alignments | AdaBoost accuracy (%) | Avg. (%) used in essential sets | Saved hours | Saved time (%) |
| default | 18.41 | 13.32 | 83.38 | 33.83 | 5.09 | 27.65 |

**Table S5:** Importance of correctly predicting the fastest inter-proteome alignments. To evaluate the importance of AdaBoost, we performed trials using a “wrong” model, which outputs the opposite prediction of the “correct” one. For example, if the model has an accuracy of 95.94% using MMseqs in the most-sensitive mode, then the “wrong” model will have an accuracy of 4.06%. Column four shows the total execution times for the homology searches using the “correct” or “wrong” AdaBoost classifier. The rightmost column shows the decrease in percentage of saved time when using the wrong model.

| Alignment tool | SonicParanoid mode | AdaBoost model | Ex. time (hours) | Saved hours | Saved time (%) | Saved time decrease (%) |
| --- | --- | --- | --- | --- | --- | --- |
| **MMseqs2** | most-sensitive | Correct | 3.91 | 2.85 | 42.18 | -70.34 |
|  |  | Wrong | 5.91 | 0.85 | 12.51 |  |
|  | sensitive | Correct | 1.65 | 0.91 | 35.60 | -58.68 |
|  |  | Wrong | 2.19 | 0.38 | 14.71 |  |
|  | default | Correct | 0.47 | 0.17 | 25.84 | -53.30 |
|  |  | Wrong | 0.56 | 0.08 | 12.07 |  |
|  | fast | Correct | 0.26 | 0.07 | 20.87 | -38.81 |
|  |  | Wrong | 0.29 | 0.04 | 12.77 |  |
| **Diamond** | most-sensitive | Correct | 1.71 | 0.63 | 27.01 | -23.11 |
|  |  | Wrong | 1.86 | 0.49 | 20.77 |  |
|  | sensitive | Correct | 0.45 | 0.17 | 27.69 | -37.29 |
|  |  | Wrong | 0.51 | 0.11 | 17.36 |  |
|  | default | Correct | 0.30 | 0.16 | 34.29 | -47.37 |
|  |  | Wrong | 0.38 | 0.08 | 18.05 |  |
|  | fast | Correct | 0.16 | 0.07 | 31.05 | -52.65 |
|  |  | Wrong | 0.19 | 0.03 | 14.70 |  |
| **BLAST** | default | Correct | 13.32 | 5.09 | 27.65 | -12.17 |
|  |  | Wrong | 13.94 | 4.47 | 24.29 |  |

**Table S6:** Saved execution time on 2,000-MAG dataset using the novel graph-based approach at default sensitivity settings. Columns two and three show the time required to perform the homology searches (in hours) using the conventional all-versus-all (complete alignments) approach and the novel approach (essential alignments), respectively. The AdaBoost accuracies and the average sizes of the original input proteins used in the essential subsets are listed in columns four and five, respectively. The last two columns on the right show the amount of time saved (in hours and percentage) when compared with the complete alignments.

| Alignment tool | Complete alignments | Essential alignments | AdaBoost accuracy (%) | Avg. (%) used in essential sets | Saved hours | Saved time (%) |
| --- | --- | --- | --- | --- | --- | --- |
| MMseqs2 | 67.38 | 56.48 | 83.89 | 28.09 | 10.90 | 16.18 |
| Diamond | 31.65 | 23.74 | 91.37 | 42.96 | 7.91 | 24.99 |

**Table S7:** Speedup folds on Broccoli and OrthoFinder when processing QfO dataset. Speedups refer to trials performed on SonicParanoid2 using only the graph-based algorithm and include the creation of ortholog groups. OrthoFinder and Broccoli used Diamond to perform the alignments, whereas SonicParanoid was executed using different alignment tools (Supplementary Table S3). Negative values represent settings for which SonicParanoid was slower.

| Alignment tool | SonicParanoid mode | Ex. time (hours) | Speed-up folds on Broccoli | Speed-up folds on OrthoFinder |
| --- | --- | --- | --- | --- |
| **MMseqs** | most-sensitive | 4.17 | -1.51 | 2.44 |
|  | sensitive | 1.91 | 1.45 | 5.33 |
|  | default | 0.75 | 3.69 | 13.57 |
|  | fast | 0.55 | 5.01 | 18.44 |
| **Diamond** | most-sensitive | 1.93 | 1.43 | 5.27 |
|  | sensitive | 0.66 | 4.17 | 15.33 |
|  | default | 0.52 | 5.32 | 19.57 |
|  | fast | 0.38 | 7.31 | 26.90 |
| **BLAST** | default | 13.67 | -4.94 | -1.34 |

**Table S8:**
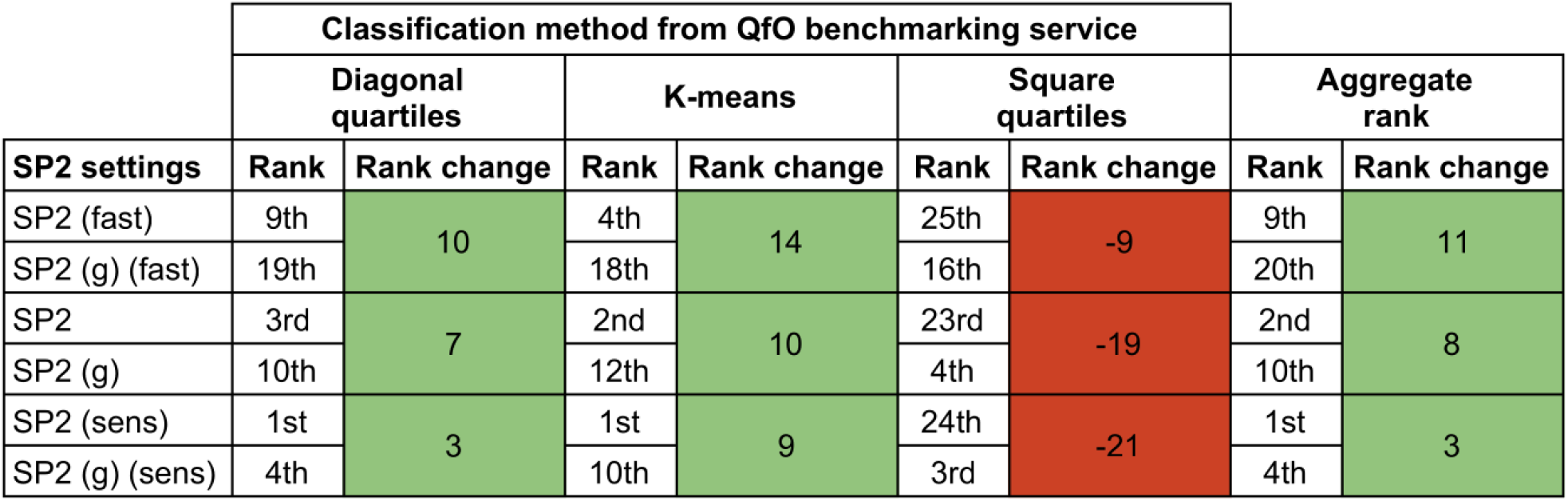
Improvements in the QfO ranking due to the inclusion of domain-based orthology. Labels containing “(g)” represent SonicParanoid2 runs in which only graph-based orthology was performed. The columns “Rank change” contain the change in number of positions resulting from the inclusion of the domain-aware algorithm. Cells with green and red backgrounds respectively represent gains and losses in the ranks.

**Table S9:**
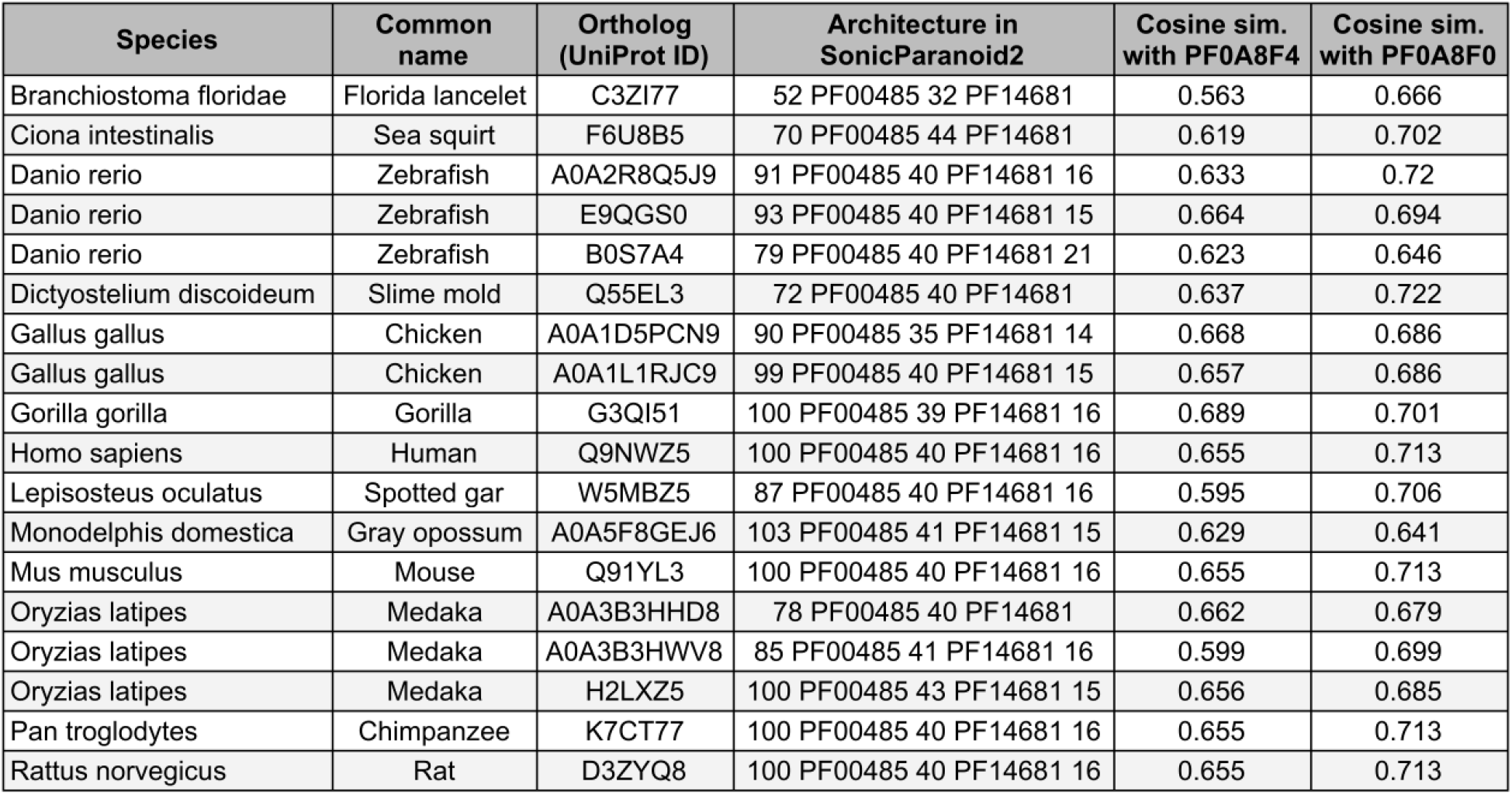
Domain-orthologs of eukaryote uridin–ecytidine kinase proteins recovered using SonicParanoid2. Third column shows a list of proteins predicted as orthologous to the human uridine–cytidine kinase proteins (Q9NWZ5), which was shown as an example of domain-fusion event by Persson *et al.* 2020. Column four shows the architectures of each protein as they appear in the training corpus for the Doc2Vec model. The last two columns show the cosine similarities assigned by Doc2Vec to each eukaryote uridine–cytidine kinase ortholog for *E. coli* PF0A8F4 and PF0A8F0 proteins, respectively.

## Equations

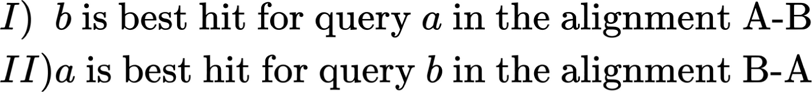

**Equation 1:** Bidirectional best hit (BBH).

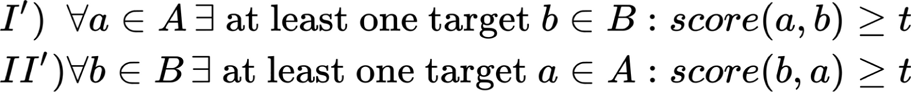

**Equation 2:** Necessary conditions for the inclusion of query and target proteins in protein subsets (*ess(A)* and *ess(B)*) used in the second inter-proteome alignment.

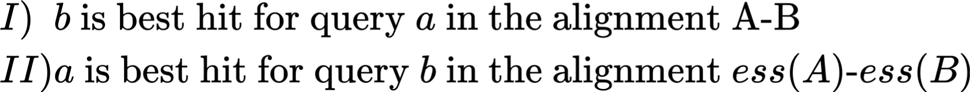

**Equation 3:** Bidirectional best hits applied to alignments conducted on essential protein subsets.

